# Forest and Biodiversity 2: a tree diversity experiment to understand the consequences of multiple dimensions of diversity and composition for long-term ecosystem function and resilience

**DOI:** 10.1101/2024.03.09.584227

**Authors:** Jeannine Cavender-Bares, Jake J. Grossman, J. Antonio Guzmán Q., Sarah E. Hobbie, Matthew A. Kaproth, Shan Kothari, Cathleen Lapadat, Rebecca Montgomery, Maria Park

## Abstract

1. We introduce a new “low-density” tree diversity experiment at the Cedar Creek Ecosystem Science Reserve in central Minnesota, USA aimed at testing long-term ecosystem consequences of tree diversity and composition. The experiment was designed to provide guidance on forest restoration efforts that will simultaneously advance carbon sequestration goals and contribute to biodiversity conservation and sustainability.
2. The new Forest and Biodiversity (FAB2) experiment uses native tree species in varying levels of species richness, phylogenetic diversity, and functional diversity to test the mechanisms and processes that contribute to forest stability and ecosystem productivity in the face of global change. FAB2 was designed and established in conjunction with a prior experiment (FAB1) in which the same set of twelve species was planted in 16 m^2^ plots at high density (0.5 m spacing). In addition to lower density plantings (1 m spacing), FAB2 also has larger plots (100 m^2^ and 400 m^2^) appropriate for testing long-term ecosystem consequences of species composition and diversity.
3. Within the first six years, mortality in the 400 m^2^ monoculture plots was significantly higher than in the 100 m^2^ plots. The highest mortality among any treatments occurred in *Tilia americana* and *Acer negundo* monocultures, but mortality for both species decreased with increasing plot diversity. These results highlight the importance of forest diversity in reducing mortality in some species and point to potential mechanisms, including light and drought stress, that cause tree mortality in vulnerable monocultures. The experiment highlights challenges to maintaining monoculture and low-diversity treatments in tree mixture experiments of large extent.
4. The FAB2 experiment provides a long-term platform for discerning the importance of species and lineage effects and of multiple dimensions of diversity in restoring ecosystem functions and services provided by forests. It also provides a platform for improving remote sensing approaches, including Uncrewed Aerial Vehicles (UAVs) equipped with LiDAR, multispectral, and hyperspectral sensors, to complement ground-based monitoring of forest function and diversity. We aim for the experiment to contribute to international efforts to monitor and manage forests in the face of global change.

## Introduction

Consequences of functional and phylogenetic diversity for ecosystem functions and services are a central topic in community and ecosystem ecology (Larkin et al. 2023, Díaz, et al. 2013), particularly in our era of rapid global change when restoring ecosystems based on sound ecological principles is critical (Leadley et al. 2022). Testing these principles based on well-designed experiments is a rapidly expanding endeavor (Paquette et al. 2018). Early forest plantation experiments in Poland, Denmark, and elsewhere demonstrated clear effects of species functional traits on ecosystem function (Vesterdal et al. 2013, Vesterdal, et al. 2008, Ladegard-Hobbie et al., 2006; Reich et al., 2005, Pedersen et al. 2005) and increasing productivity with functional diversity in forest composition experiments in Costa Rica (Ewel et al., 2015; Haggar & Ewel, 1997). These experiments were consistent with species effects in herbaceous and shrub ecosystems (Hobbie, 1994, 1995, Wedin & Tilman 1990) and dovetailed with early diversity-ecosystem function experiments in herbaceous systems, including the BioDIV experiment at Cedar Creek Ecosystem Science Reserve (Tilman, 1993, 1994) and the distributed diversity experiments in Europe, centered in Jena (Eisenhauer et al., 2019, Hector et al. 1999). The herbaceous experiments, in particular, have demonstrated that above ground processes, including those that can be remotely sensed, are linked to and predict belowground processes (Cavender-Bares et al., 2021; Cline et al., 2018). The insights provided by these studies led to calls for a tree diversity network to determine whether diversity in forests in contrasting ecosystems would result in consistent enhancements of productivity as in early experiments in forests and grasslands (Verheyen et al., 2016). Experiments designed to test the effects of tree composition and diversity on ecosystem function and resilience have now emerged across the globe, led by TreeDivNet and the IDENT experiments (Paquette et al., 2018).

Results from the new masting of forest diversity experiments have upheld previous findings that greater diversity contributes to higher productivity through a combination of complementarity and species effects (Grossman et al., 2017; Tobner et al., 2016; Williams et al., 2017). In addition, these new experiments have expanded on prior findings, including showing that greater multifunctionality translates to different kinds of ecosystem services (Messier et al., 2021) and increased resilience through reduced heterogeneity in survival and growth of forest stands in response to drought (Hutchison et al., 2018). These experiments are critical for guiding forest restoration practices (Grossman et al., 2018), given efforts linked to the Kunming Montreal Global Biodiversity Framework as part of the UN Convention on Biodiversity to provide guidelines and criteria for restoring degraded ecosystems (Leadley et al., 2022). They also provide a platform for integrating remote sensing tools with measured biological processes on the ground and for advancing capabilities to monitor forest growth, diversity and ecosystem function (Cavender-Bares et al., 2022; Williams et al., 2021; Cavender-Bares et al., 2020). Such tools are urgently called for to advance the global monitoring of biodiversity and ecosystem function (Gonzalez et al., 2023).

### The Forest and Biodiversity (FAB1) high density experiment

Our team participated in the first phase of these experiments by designing and establishing the Forest and Biodiversity (FAB) 1 experiment at the Cedar Creek Ecosystem Science Reserve in 2013 as part of IDENT (Grossman et al., 2017). FAB1 includes 12 native tree species in a combination of monocultures, two-species, five-species and 12-species plots. The trees were planted at high density (0.5 m apart) in small plots (16 m^2^). The experiment was designed in part to test for distinct effects of phylogenetic and functional diversity, recognizing that evolutionary history influences species traits and interactions and that phylogenetic and functional dimensions of biodiversity are not always congruent (Cavender-Bares et al., 2009). We thus build on studies with herbaceous species that show that shared ancestry (Cadotte et al., 2008; Cadotte & Dinnage, 2012) and functional diversity (Cadotte et al., 2009; Flynn et al., 2011) influence diversity-ecosystem function relationships beyond species richness.

The FAB1 experiment has been primarily used to test the consequences of tree diversity and composition for 1) ecosystem functions, including aboveground productivity (Grossman et al. 2017) and soil processes (Maillard et al. 2023), 2) the composition and abundance of other trophic levels (Grossman et al. 2019ab) and 3) the mechanisms underlying these effects (Kothari et al. 2021). Importantly, FAB1 was one of the first of the new wave of forest diversity experiments to provide evidence for overyielding, meaning that on average, diverse mixtures had greater productivity than expected based on monoculture productivity (Bryant et al., in review; Grossman et al., 2017). Most of the overyielding could be attributed statistically to positive complementarity effects (Loreau & Hector, 2001), and most plots had weak or negative selection effects. Diversity both increased and alleviated the impact of enemy-mediated biotic feedbacks caused by fungi, bacteria, and invertebrate animal pests (Grossman et al., 2019). The mechanisms underlying these ambivalent responses of natural enemies to diversity are also likely to be heterogeneous. For instance, it has been proposed that, in polycultures, individuals can vary considerably in height relative to their neighbors; some become more apparent to herbivores and therefore experience more damage (Damien et al., 2016; Endara & Coley, 2011). In FAB1, we found this to be the case for two broadleaf species: *Acer negundo* and *Betula papyrifera*. But we found that, for four *Quercus* spp. and *Tilia americana*, more apparent individuals (those that were much taller than their neighbors) actually experienced less herbivory (Grossman et al., 2019). In this case, diversity can have complex consequences for natural enemy damage via even a single mechanism, such as plant apparency.

Complementarity can be driven by multiple mutually inclusive mechanisms, including resource partitioning, abiotic facilitation, and biotic feedbacks mediated by shared enemies or mutualists (Barry et al., 2019). Research in FAB1 tested multiple of these mechanisms and showed that abiotic facilitation by fast-growing trees (mostly conifers) aided the growth of the most shade-tolerant species in the experiment (Kothari et al., 2021). There was indirect evidence for resource partitioning based on the association between increased growth rates, light interception and complementarity in high-dimensional spectral phenotypes (Schweiger et al., 2021). Thus, FAB1 provided evidence for multiple distinct mechanisms of complementarity and at least indirect evidence for resource partitioning.

In FAB1, diversity (including taxonomic, functional and phylogenetic diversity) also influenced ecosystem processes in ways that could cause feedbacks for productivity. Diverse litter mixtures had slower decomposition rates, particularly for labile carbon pools (Grossman et al., 2020) and plot-level differences in tree diversity had already begun to shape the soil microbial community within three years after experimental establishment (Grossman et al., 2019; Maillard et al., 2023). There was some evidence that species diversity increased soil carbon sequestration, but soil carbon was surprisingly unrelated to aboveground productivity after seven years of growth (Bryant et al., in review).

The high-density of FAB1 was designed to promote interactions among neighboring trees relatively quickly, allowing us to detect diversity effects and probe their mechanisms at an early stage. However, inferences based on experiments with very high stem density may not be representative of natural tree communities, which show a wide range of stem densities, or forest plantations, which are often much sparser (Gabira et al. 2023). Moreover, the small plot size of FAB1 could make it hard to identify treatment effects, particularly on ecosystem functions that have large spatial footprints and may bleed between adjacent plots. These concerns motivated the establishment of a lower-density Forest and Biodiversity experiment, FAB2. A larger plot size and expanded experimental design also lend themselves well to the rapidly increasing use of remote sensing tools for assessing aboveground community dynamics and ecosystem functions. Here, we document the FAB2 experimental design (Figure 1) and rationale, present the hypotheses we sought to test (Box 1), and describe the site management, initial planting, mortality and adjustments made in response. We include a series of definitions (Box 2) to help clarify our hypotheses and conceptualization of what the experiment was designed to test.

**Figure 1.**
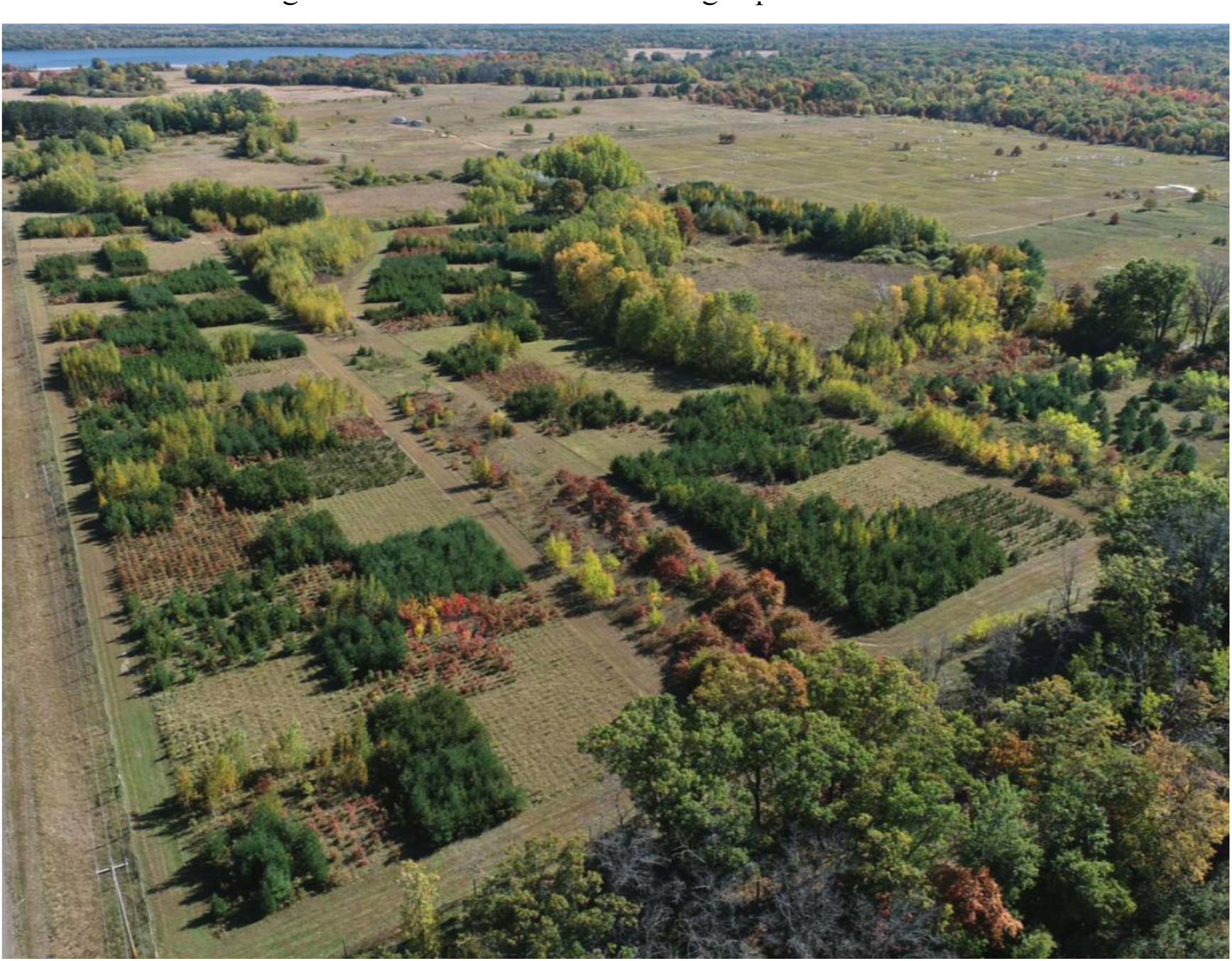
An aerial view of the forest and biodiversity (FAB2) experiment in October 2023 at the Cedar Creek Ecosystem Science Reserve, showing the combination of 100 m^2^ and 400m^2^ plots. Differences in color and size of evergreen conifers and deciduous angiosperms are evident.

### Goals of the FAB2 experiment

We designed the FAB2 experiment (Figure 1) to determine the influence of tree diversity and composition on long-term community processes and ecosystem function and resilience. Our main objective is to experimentally test hypotheses that generate informed insights to aid forest restoration. We are specifically interested in understanding the consequences for other trophic levels and for ecosystem functions of 1) tree species and lineages and 2) tree diversity; and in 3) deciphering the mechanisms by which tree species influence each other with consequences for ecosystem processes. Consequently, we are examining influences of tree composition and diversity on the abiotic environment including resource availability and stress, as well as cascading impacts on biotic processes, including herbivory and microbial processes. Our central hypotheses are posited in Box 1.

FAB2 differs from other tree diversity experiments in having larger plot sizes than most and including very large-sized monocultures to test specifically for species and lineage effects on ecosystem processes. It is also designed to test the importance of functional and phylogenetic diversity within a given species richness level. Given the large size of the experiment, we are developing remote sensing approaches for cost-effective long-term measurement of growth, survival, structure, phenology, and ecosystem processes. Our aim is to use relatively inexpensive (multispectral, thermal, LiDAR) and advanced (hyperspectral) sensors on UAV platforms to characterize phenological changes in forest structure and canopy chemistry to test our hypotheses. We are using this remotely sensed information to model multiple dimensions of biodiversity as well as ecosystem productivity and will relate them to belowground processes such as soil nutrient availability and soil processes. We anticipate variability among assemblages in when canopy closure occurs with consequences for the timing and rate of influence on soil processes.

## Methods

### The Forest and Biodiversity (FAB2) low density experiment

In 2016, we established the first phase of a low-density tree experiment (FAB2). We planted trees 1 m apart in large plots (100 m^2^), with replicated monocultures and two-species, four-species, six-species and twelve-species polycultures (Table 1). In 2017, we planted each species in replicated monocultures and in five 12-species polycultures at 400 m^2^. The full experiment covers approximately 6.5 hectares (Figure 1). FAB2 was established in an abandoned old field dominated by herbaceous species at the Cedar Creek Ecosystem Science Reserve, a 2300 ha reserve and National Science Foundation Long Term Ecological Research site in eastern Minnesota, USA (45°25’ N, 93°10’ W). The site is situated on excessively drained sandy soils (Alfic Udipsamments, Grigal, 1974) of the Anoka Sand Plain and has a humid continental climate with warm summers and cold winters. Cedar Creek is located at the boundary of the Midwestern tallgrass prairies, Eastern deciduous forests, and Northern boreal forests. In the year prior to planting, the experimental site was treated by removing nearby trees and stumps that could cast shade, tilling the topsoil and burning the herbaceous vegetation. The experiment was fenced to exclude large mammalian herbivores. We mowed between rows of planted trees and occasionally hand-weeded 2-3 times during the growing season each year during the first three years, and at least once per year in subsequent years to minimize competition from herbaceous vegetation. Trees were fully irrigated in spring and summer in the first six years of the experiment (2016-2022) using two Kifco T200L water reels. In 2023, trees were irrigated using sprayers only during periods with lower precipitation than average. Once canopy closure occurs, trees are no longer irrigated, a point which has already occurred for many assemblages. From 2016-2022, *Spermophilus tridecemlineatus* (gophers) were monitored and trapped every year to minimize disturbance.

**Table 1.** General design of FAB2 with the number of plots in each species richness level, the description of the treatments and how they were selected, and the size of the plots. There were originally 400 m2 monoculture plots but only 25 of those remain, due to mortality. The Oak-DIV experiment, nested within the larger experiment, includes 30 plots in total, including three monocultures of each of the four oak species plus three replicates of all of the two and four species combinations. Some of the Oak-DIV plots are also part of the general FAB2 experiment.

| No. plots | Species richness | Description | Size |
| --- | --- | --- | --- |
| 36 | 1 | 12 monocultures replicated 3x | 100 m<sup>2</sup> |
| 47 | 2 | 11 random; 36 selected for different PV-FV combinations | 100 m<sup>2</sup> |
| 45 | 4 | 9 random; 36 selected for different PV-FV combinations | 100 m<sup>2</sup> |
| 10 | 6 | Random | 100 m<sup>2</sup> |
| 10 | 12 | All species polycultures | 100 m<sup>2</sup> |
| 36* (25) | 1 | Monocultures | 400 m<sup>2</sup> |
| 5 | 12 | All species polycultures | 400 m<sup>2</sup> |
| 30 | 1, 2, 4 | Oak-DIV | 100 m<sup>2</sup> |

### Tree diversity gradient across functional and phylogenetic variability levels in 100 m^2^ plots

The main part of the FAB2 experiment includes 148 plots each of 10 x 10 m (100 m^2^) planted in species richness levels of 1, 2, 4, 6 or 12. Of these, 36 plots were monocultures (twelve species each replicated in three monocultures), 47 plots were bicultures, 45 plots were four-species mixtures, ten plots were six-species mixtures and ten plots were twelve-species mixtures. For the bicultures, 11 were combinations of randomly selected species and 36 were bicultures of species selected to achieve targeted phylogenetic variability (PV) – functional variability (FV) combinations, also calculated in terms of phylogenetic diversity (PD) – functional diversity (FD) (details below). For the four-species mixtures, nine were combinations of randomly selected species, and 36 were mixtures of species selected based on achieving targeted PV–FV combinations. The six-species mixtures were all created through random selection of species. Survival was approximately 95% per year on average across all species (Table 3). All seedlings that died were replanted each year between 2017-2020.

### Selection of native tree species

Prior to designing both FAB experiments, we planted 22 native tree species into an old field at CCESR without watering or other amendments. For FAB1 and FAB2, we chose only species that had survival rates higher than 50% in the initial planting. Our final planting list of 12 native species for the FAB experiments included eight angiosperms, including four oak species (*Q. alba, Q. ellipsoidalis*, *Q. macrocarpa* and *Q. rubra*), a birch (*B. papyrifera*), two maple species (*A. rubrum* and *A. negundo*), and basswood (*Tilia americana*), as well as four conifers, including three pines (*Pinus banksiana, P. resinosa* and *P. strobus*) and eastern red cedar (*Juniperus virginiana*).

### Tracking maternal families for five species

To track genetic identities within populations and account for genetic variation within some of the species, individuals from half-sib families were planted for five species—*Acer rubrum, Betula papyrifera, Quercus ellipsoidalis, Q. macrocarpa* and *Q. rubra.* These individuals were randomly assigned to different treatments (Table 2). Seeds were collected from known and geolocated mother trees in Minnesota (Figure S1), germinated and grown in the PRT Dryden nursery in Dryden, ON, Canada (www.prt.com), and returned to Minnesota after one year of growth. Individuals from known families (based on mother identities) were randomly assigned to different treatments in which those species were represented.

**Table 2.** Scientific and common names of species and taxonomic families included in FAB2. The number of maternal families raised for inclusion and tracking in the 100 m^2^ experimental plots are shown for each of the five tree species for which we collected our own seed stock. The number of monocultures for each species for the two plot sizes is shown.

| Species | Family | Common name | Maternal lines tracked | 100 m<sup>2</sup> monocultures | 400 m<sup>2</sup> monocultures |
| --- | --- | --- | --- | --- | --- |
| *Acer negundo* | Sapindaceae | Box elder |  | 3 | 3 |
| *Acer rubrum* | Sapindaceae | Red maple | 20 | 3 | 3 |
| *Betula papyrifera* | Betulaceae | Paper birch | 21 | 3 | 0 |
| *Juniperus virginiana* | Cupressaceae | Red cedar |  | 3 | 3 |
| *Pinus banksiana* | Pinaceae | Jack pine |  | 3 | 3 |
| *Pinus strobus* | Pinaceae | White pine |  | 3 | 3 |
| *Pinus resinosa* | Pinaceae | Red pine |  | 3 | 3 |
| *Quercus alba* | Fagaceae | White oak |  | 3 | 1 |
| *Quercus ellipsoidalis* | Fagaceae | Northern pin oak | 13 | 3 | 3 |
| *Quercus macrocarpa* | Fagaceae | Bur oak | 12 | 3 | 3 |
| *Quercus rubra* | Fagaceae | Red oak | 18 | 3 | 1 |
| *Tilia americana* | Malvaceae | American basswood |  | 5 (3x100 m<sup>2</sup> + ½ of 400 m<sup>2</sup> plot) | 0 |

### Large 400 m^2^ monocultures and polycultures

Large monocultures and twelve-species polycultures in 20 x 20 m plots (400 m^2^) were planted in an arrangement that dispersed them throughout the experimental area, again with 1 m spacing between trees (Figure 1). The rationale for these larger plot sizes was to reduce edge effects, blown-in litter and root in-growth from neighboring assemblages to test for long-term ecosystem effects that might not emerge in smaller plots. Originally, all twelve species were planted in large monocultures replicated 3x along with five twelve-species polycultures. In spring 2022, due to low initial survival for *Tilia americana*, *Acer negundo, Quercus ellipsoidalis, Q. alba* and *Betula papyrifera* in 400 m^2^ monocultures, eleven 400 m^2^ monoculture plots were removed from the experiment (Table 2) and converted to 10 x10 m plots to create the Oak-DIV experiment that compares the interspecific interactions of oak species from the same and different lineages (*Quercus* section *Quercus*, white oaks vs *Quercus* section *Lobatae*, red oaks). No large monocultures remained for *T. americana.* One large monoculture was retained for *B. papyrifera*, *Q. alba* and *Q. rubra,* and two for *A. negundo* (Table 2).

### Phylogenetic and functional variability and diversity

The mixture of close relatives and distant relatives with a range of trait combinations allowed us to create mixtures of (1) low phylogenetic and functional variability, (2) high phylogenetic and functional variability, (3) low phylogenetic and high functional variability, and (4) high phylogenetic and low functional variability. These combinations were previously tested in two and five species mixtures in FAB1 (Grossman et al., 2017) and the long-term consequences of these interactions are still being tested (Bryant et al., in review). In the FAB2 experiment, we created more combinations by creating two-, four- and six-species mixtures (Table 1).

During the design phase, phylogenetic variability and diversity were calculated based on the phylogenetic tree published by Zanne et al. (2014) with modifications. We calculated phylogenetic species variability (Helmus, 2007) in Picante (Kembel et al., 2010), which we call phylogenetic variability (PV) for simplicity, and phylogenetic diversity (PD) based on Faith’s PD. PV varies between 0 and 1 and is independent of species richness. After planting, we recalculated these metrics using the Smith and Brown 2018 v.01 megaphylogeny (Smith & Brown, 2018), pruned to the 12 FAB species and ultrametricized using phytools (Revell, 2012). Both pruned trees are shown in Fig. S3.

We calculated functional variability (FV) and functional diversity (FD) from the same metrics using a multivariate trait dendrogram treated as a bifurcating phylogeny (Cadotte et al. 2009, Cavender-Bares and Reich 2012). Each trait was scaled with a mean of 0 and SD of 1. Using the hclust algorithm in R, we created a cophenetic distance matrix and the trait dendrogram. We chose this method of calculating functional diversity to be directly comparable with the method of calculating phylogenetic diversity. During the design phase, functional variability and diversity were based on species-level measured values of leaf mass per unit area (LMA) and leaf nitrogen concentration (N_mass_) from the GLOPNET database (Wright et al., 2004), ranked values of wood density, shade tolerance and drought tolerance based on the authors’ expert knowledge of the trees, and binary values of leaf habit (evergreen/deciduous), mycorrhizal type (ECM or AM) and calcium use (high/low). We recalculated functional variability and diversity here using measured values of LMA, N_mass_ (Sendall & Reich, 2013; Wright et al., 2004) and wood density (Jenkins et al., 2003), binary values of leaf habit, mycorrhizal type (Averill et al., 2019) and calcium use, and indices of shade and drought tolerances (Niinemets & Valladares, 2006). We chose these traits to encompass functions related to nutrient use, including micronutrients important for soil processes and soil organisms, water use and drought tolerance, and light use and shade tolerance. These traits are linked to the major axes of environmental variation important for niche partitioning in natural forest communities of the region.

Within each species richness level, PV was binned into eight quantiles and compared to FV. Species pairs or four-species mixtures were randomly drawn from each quantile in a manner that would create low PV-high FV, low FV-high FV, high PV-high FV, and low PV-low FV combinations, to the extent possible. The range of variation on these two axes is shown in Figure S2 for the four-species mixtures. We also recalculated PD and FD (rooted and unrooted, Faith, 1992) using the updated input data described above (supplemental information). The spatial arrangement of PV and FV plots is visualized in Figure 3.

## Data analysis

We fit linear models to the tree mortality data to test for the effects of plot diversity and composition, species and plot size. Percent mortality was calculated for each species in each plot where it occurred for every year. We ran all models as untransformed percent mortality as well as arcsine transformed values. In the first set of analyses using only 100 m^2^ plots, percent mortality was treated as the independent variable. Species, plot diversity and their interaction were the predictor variables. In five separate analyses, species richness, functional diversity, functional variability, phylogenetic diversity or phylogenetic variability were treated as the diversity variable. In a second set of analyses, using only 400 m^2^ plots, percent mortality was treated as the independent variable predicted by species, plot diversity and their interaction. Again, species richness, phylogenetic variability, phylogenetic diversity, functional variability, or functional diversity were treated as the diversity variable. In a third set of analyses, both 100 m^2^ and 400 m^2^ plots were included. Percent mortality was treated as the independent variable, and the predictor variables were plot size, species and plot diversity, as well as the interactions of species with plot size, and plot diversity. Finally, a last analysis using both 100 m^2^ and 400 m^2^ plots treated percent mortality as the independent variable and treated as predictor variables species, plot size, the proportion of angiosperms in the plot and the interactions of species with plot size and with proportion of angiosperms in the plot. There was no substantial difference in results when percent mortality or arcsine transformed mortality values were used, indicating that the model results are robust to normality assumptions. Model results using transformed data are reported in the text, but untransformed data are used for the figures. Linear models were run in JMP version 16.0 (SAS Institute Inc., Cary, NC, 1989–2023).

### Box 1 Conceptual figure and hypotheses

FAB2 (Figure 2) seeks to test the following hypotheses about the consequences of i) **species and lineages** and ii) **taxonomic, functional and phylogenetic diversity and community composition** for ecosystem functions and other trophic levels as well as iii) the underlying **causal mechanisms**. Our central hypotheses in these realms are posited here.

**Figure 2.**
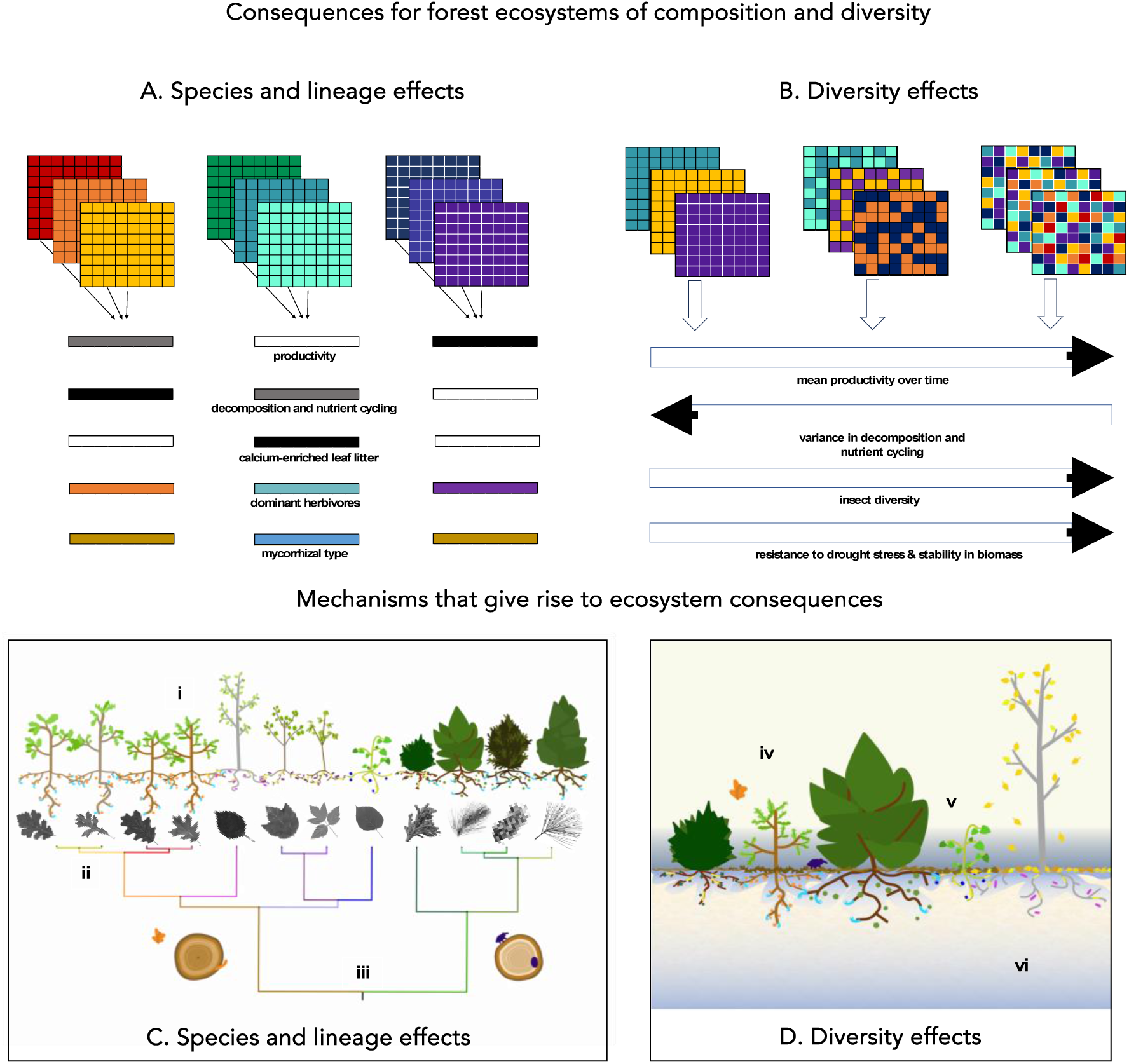
The forest and biodiversity (FAB2) experiment is designed to test consequences of tree species and lineages and of tree diversity for ecosystem functions and other trophic levels and to uncover the mechanisms underlying these effects. *Clockwise from top left:* **A** shows the consequences of species and lineage effects of closely related monocultures in three lineage groups. Closely related species will exhibit similar patterns in productivity, decomposition and nutrient cycling, calcium-enriched leaf litter, dominant herbivores, and mycorrhizal type. **B** Examples of the consequences of tree diversity treatments ranging from varied monocultures (left) to bicultures of different phylogenetic and functional similarity (middle) and higher diversity treatments (right). With greater tree diversity, mean productivity is expected to increase, variance in decomposition and nutrient cycling is expected to decrease, local insect diversity is expected to increase and resistance to drought and the stability of biomass in forest ecosystems are expected to increase. **C** shows the mechanisms underlying species and lineage effects associated with the experimental design. Included in the tree diversity experiment are twelve tree species native to Minnesota that span a wide range of phylogenetic and functional diversity. From left to right: *Quercus macrocarpa, Q. alba, Q. rubra, Q. ellipsoidalis, Betula papyrifera, Acer rubrum, A. negundo, Juniperus virginiana, Pinus resinosa, P. banksiana, P. strobus.* The experimental design enables the study of **(i)** plant functional traits and intrinsic growth rates, **(ii)** host specificity and co-evolutionary acquisition of symbionts, including bacterial and fungal partners, and **(iii)** the deep evolutionary divergence in wood and leaf structural properties and their consequences for ecosystem processes. **D** depicts the mechanisms underlying tree diversity effects that can be studied in the experiment. These effects include **(iv)** dilution effects, **(v)** phenological offsetting in light and nutrient use, and **(vi)** facilitation through shading and soil moisture maintenance. Artwork in panels C and D was created by Maria Park.

**Figure 3.**
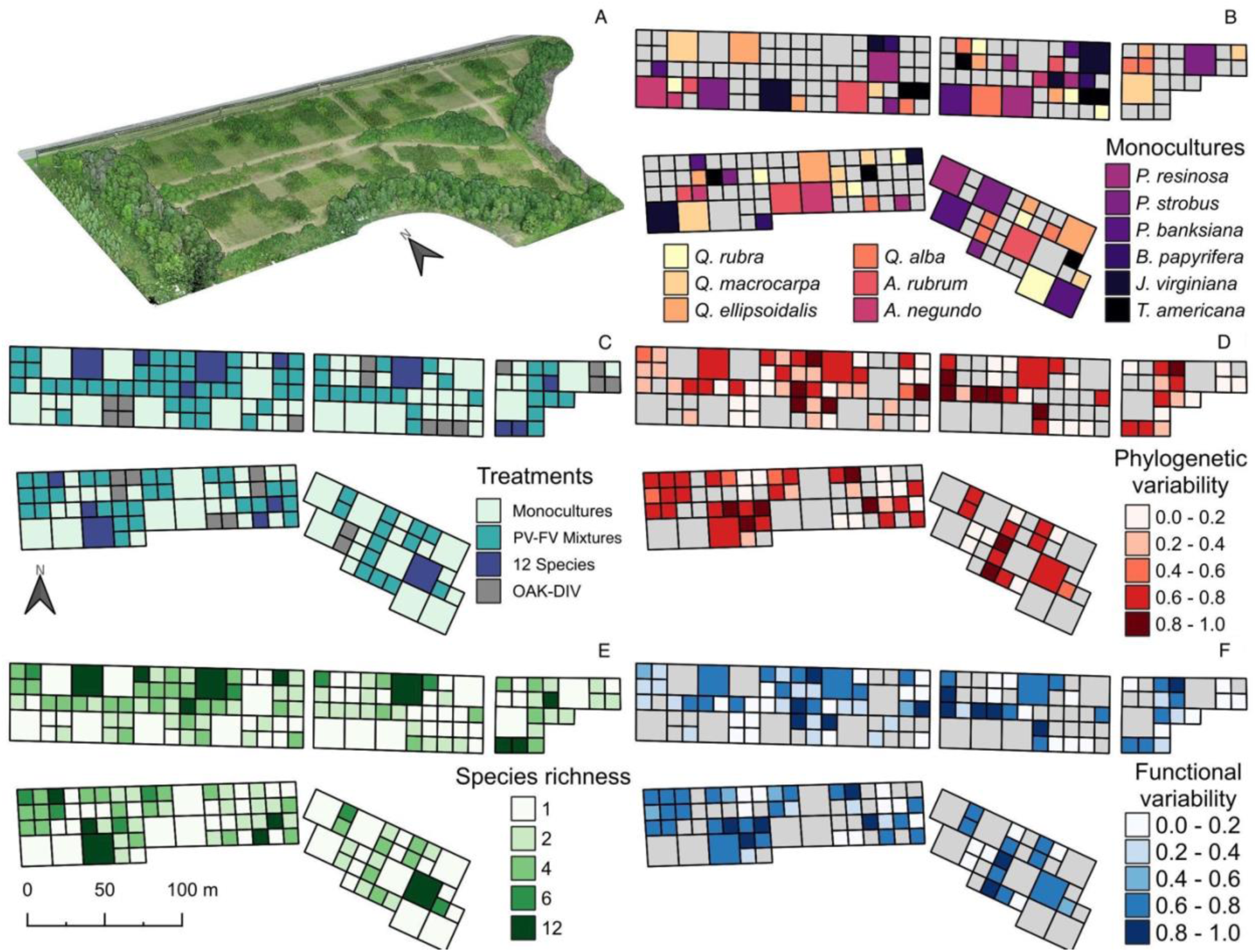
Experimental design of the Forest and Biodiversity Experiment 2 at the Cedar Creek Ecosystem Science Reserve. A) A LiDAR image during summer 2022. Color images show the spatial arrangement of B) species composition of plots with monocultures, C) monocultures, PV-FV mixtures, 12 species mixtures and the oak species mixtures within the Oak-DIV nested experiment, D) Phylogenetic variability within plots, E) Number of species in each plot, and F) phylogenetic variability within plots.

#### Species and lineage effects

**Productivity.** Ecosystem function in monocultures will differ among plots planted with species or lineages that have contrasting functional trait values, particularly for “effect traits” (*sensu* Lavorel & Garnier, 2002) known to have ecosystem consequences. For example, we expect young monocultures to vary in productivity due to a range of growth strategies among species and lineages.

**Decomposition and Nutrient Cycling.** We also expect divergence in nutrient cycling due to contrasting properties of leaf and root detritus and exudates as well as mycorrhizal types (Phillips et al., 2013). Due to shared ancestry that influences trait expression, effect traits will cause a phylogenetic signal in some ecosystem functions. Species and lineage effects on ecosystems resulting from chemical traits that influence soil processes are expected to become stronger through time. These effects will be stronger with increasing plot size due to reduced influence from neighboring assemblages.

**Soil Community.** Phylogenetically distinct monocultures will diverge in fungal and bacterial soil communities more than phylogenetically similar monocultures. This hypothesis is based on the postulate that shared ancestry results in similar decomposing root and leaf litter properties and similar fungal symbionts and soil pathogen communities as a consequence of host-specificity. These effects will become stronger through time.

**Natural Enemies.** Phylogenetically distinct monocultures will diverge in specialist herbivores, pathogens and enemy community compositions due to host specificity. These effects will become stronger as plot biomass increases through time and will increase with plot size due to greater host species apparency.

**Mycorrhizal Type and Genotypic Effects.** Fungal root and endophytic symbionts as well as specialist herbivores will show composition effects of host genotype within a species due to host specificity.

#### Diversity and composition effects

**Productivity above and below ground.** In general, greater numbers of species, which on average will covary with greater phylogenetic and functional diversity, will result in greater forest productivity due to complementarity effects that result in greater overall resource use, dilution of natural enemies, and/or facilitation (including amelioration of physical stress). At the same time, highly productive species will contribute to high ecosystem productivity where they occur due to sampling (“selection”) effects. Both effects will contribute to belowground processes including soil carbon accumulation. More functionally, phylogenetically, spectrally, and taxonomically diverse plots will have greater overyielding in soil carbon accumulation and *greater overall soil carbon accumulation* because of their higher productivity.

**Complementarity vs. selection in relation to similarity.**

We expect that assemblages with more phylogenetically, spectrally and functionally similar species— both within a species richness level as well as across all richness levels—will show less niche partitioning, structural diversity and facilitation, and therefore lower complementarity effects than those with more distantly related, functionally dissimilar and spectrally dissimilar species. We thus expect mixtures with more distantly related, functionally dissimilar and phenotypically dissimilar species to result in greater net biodiversity effects (overyielding). We expect multiple mechanisms of complementarity may be involved related to functional differences in stress response and resource use as well as host specificity. At the same time, we also expect high variability in productivity across monocultures and two species assemblages, due to a range of growth strategies. “Selection” (or sampling) effects are anticipated, whereby highly productive species contribute disproportionately to total biomass of a mixture, particularly for the conifers in the system. Over time, disease and other stressors may reduce variation. Both complementarity effects and “selection” effects on productivity are expected to become amplified through time.

**Facilitation vs. Competition.** We hypothesize that species that ameliorate stress of neighbors through temperature/microclimate and light impacts can increase growth, productivity, and survival rates of neighbors. Species that are susceptible to high light, low water, low nutrient or temperature stress are more likely to benefit from facilitation by their neighbors, while species that are less susceptible to such stress factors are more likely to face competition. Survival and growth are likely to be impacted more by facilitation effects in young individuals than established individuals.

**Trophic Consequences of Productivity.** Due to greater productivity, diverse plots will support more abundant *generalist* herbivore communities compared to less productive assemblages. Both within a species richness level and across the experiment, there will be a positive relationship between the tree assemblage phylogenetic, functional and phenotypic diversity and the mass of mostly generalist arthropod consumers (leaf chewers and skeletonizers). This relationship is expected due to combined effects of greater plant productivity from net biodiversity effects that support greater secondary production, as well as greater host diversity that support more diverse herbivore communities, including herbivore taxa of higher mass.

**Trophic Consequences of Diversity.** More diverse assemblages of primary producers are expected to support more diverse communities across trophic levels because more diverse niches will be available for consumers. Influences of tree assemblages for soil and aboveground community partners will depend on both composition (species and lineage effects) and diversity effects. With increasing tree diversity, especially phylogenetic diversity, in assemblages, *specialist* consumers (foliar fungi as well as galling and leaf mining arthropods) will decline in abundance (galls) or severity (fungi), because of dilution effects (Eisenhauer et al., 2019; Muiruri et al., 2019) but will increase in diversity with increasing tree diversity. Similarly, soil community partners, including fungal root and endophytic symbionts, will show composition effects of both host identity as well as neighboring species identities given the genetically-programmed interactions between them. Diverse tree mixtures, particularly more phylogenetically diverse mixtures, will provide more diverse niches for soil inhabitants and result in higher microbial diversity. Over time, diverse mixtures will tend to converge in microbial diversity and composition due to host specificity of fungal symbionts and soil pathogens and on functional attributes that influence soil inputs (soil and exudates).

**Stability, Resistance and Resilience.** We expect more diverse mixtures, those with a range of heights, foliar drought resistance and rooting depths, for example, to show greater resistance to chronic and episodic drought and greater stability of stem and height growth through time. We thus anticipate that resistance (integrated stability), including the resistance to and recovery of ecosystem properties in response to a disturbance and climatic stress (Helfgott, 2015; Yi & Jackson, 2021) and resilience (the ability of ecosystem properties to return to a pre-disturbance condition) after a disturbance (Fraccascia et al., 2018; Yi & Jackson, 2021) will be higher in diverse mixtures than monocultures.

**Long-term Dynamics.** We expect that the effects of initial diversity treatments will tend to compound over time and affect long-term dynamics in the fitness of particular plants, influencing the composition and stability of mixtures.

**Fitness.** At the individual tree level, fitness, as measured by joint survival, growth and fecundity, will be higher on average in multiple species mixtures than in monocultures, given net fitness benefits of greater resource partitioning and lower stress for individual trees. However, over time, overtopping may result in lower fecundity of subordinates, and self-thinning may result in loss of fitness of non-dominant individuals.

**Dominance.** In multiple species mixtures, species dominance within plots will vary with composition. Conifers will become dominant in plots if initially planted, due to their fast growth and shading (high leaf area index). Angiosperms will only become dominant in mixtures where conifers are excluded. In mixtures that ultimately become dominated by particular species, species or lineage consequences for ecosystem function will be more important than diversity effects.

**Succession.** Over decadal time scales, differential mortality across species mixtures will result in thinning and changes to community composition over time (“succession”). Due to competition and facilitation effects described above, assemblages planted with high functional diversity (FV, FD) will undergo composition changes, initially becoming more similar (decreasing in FV, FD), then slowly becoming more dissimilar before becoming more similar again (as early-successional species are replaced).

### Box 2 Definitions

#### Species and lineage effects on ecosystem processes

Evolved differences among species and lineages can alter ecosystem processes through influences on resource supply, disturbance, biogeochemical processes, biophysical processes, or trophic interactions (Chapin et al., 2011; Hobbie, 1995, 2015). Greater variation in ecosystem processes expected among lineages than among closely related species that have more shared ancestry.

#### Diversity components

Multiple dimensions or components of biodiversity capture different aspects of ecological variation, including taxonomic diversity, phylogenetic diversity, functional diversity and spectral diversity. Taxonomic diversity focuses on the diversity of species or between higher-order clades (Simberloff, 1970); phylogenetic diversity measures the evolutionary distances between species or individuals since divergence from a common ancestor (Faith, 1992); functional diversity (e.g., Lavorel et al., 2008; Mouchet et al., 2010; Villeger et al., 2008) quantifies variation among species as a consequence of measured differences in their functional traits; spectral diversity (Dahlin, 2016; White et al., 2010) measures the variability in spectral reflectance from vegetation (or from other surfaces), either measured and calculated among individual plants or, more commonly, calculated among pixels or among other meaningful spatial units (Cavender-Bares et al., 2020).

**Community composition** The identity (including taxonomic, functional, phylogenetic or spectral, etc.) and relative abundance of different species, lineages or groups within a particular ecological community or ecosystem.

#### Apparency

The likelihood that an animal or microbe can encounter its plant partner. The degree to which a focal plant is apparent depends not only on the intrinsic qualities of the plant, but also on the physical structure of surrounding vegetation (Endara and Coley, 2011; Castagneyrol et al., 2013). For instance, damage from pine processionary moth, a specialist pine herbivore, is ameliorated when pines are interspersed with fast-growing birch neighbors, which make pines less “apparent” to their would-be herbivore (Damien et al. 2016).

#### Complementarity

Interactions of species that result in greater use of limiting resources (Hooper & Dukes, 2004; Williams et al., 2017) or reduced biotic or abiotic stress. Complementarity effects can be measured by the ecosystem performance of mixtures relative to that of monocultures (Loreau & Hector, 2001).

#### Dilution effects

The resource concentration hypothesis (Root, 1973; Tahvanainen & Root, 1972) assumes that specialist natural enemies will become more locally abundant and therefore cause more damage to host plants when their hosts are locally abundant. Following from this assumption, the “dilution” of preferred hosts through mixing with non-hosts in a diverse polyculture will reduce the local abundance/prevalence of specialist natural enemies, leading to lower levels of damage to hosts (Keesing & Ostfeld, 2021).

#### Dominance

When neighboring plants compete for resources, their access to these resources may be proportionate to plant size (symmetric competition) or larger plants may have disproportionate access (asymmetric competition (Schwinning & Weiner, 1998; Fernández-Tschieder & Binkley, 2018). The latter case is often referred to as dominance. In cases of extreme competitive asymmetry, dominant individuals may monopolize access to key resources such as light or water, severely reducing the growth of subordinate neighbors and eventually killing them, reducing the positive relationship between biodiversity and ecosystem functioning when dominant species are present (Crawford et al., 2021).

#### Ecosystem functions or ecosystem functioning

The sizes of pools of materials or energy (e.g., carbon, nitrogen, or biomass) and rates of processes, including fluxes of materials or energy. High or low values are not inherently good or bad (Hooper et al., 2005).

#### Facilitation

Interactions in which a given individual or species shows greater performance due to the presence or increasing density of other individuals or species in its vicinity. In plant ecology, facilitation is usually taken to include only non-trophic interactions between independent plants (Brooker et al., 2008); however, it may be mediated by those plants’ interactions with natural enemies or symbionts, alongside various abiotic mechanisms (Wright et al., 2017). In a pairwise context, facilitation is often discussed in terms of the positive effects of “benefactor” species on “beneficiary” species.

#### Host specificity

A phenomenon in which an organism of one trophic level has evolved to depend solely on an organism of another trophic level or its close relatives.

#### Niche partitioning

Tightly related to resource partitioning. The process by which natural selection drives competing species into different niches and patterns of resource use, thereby reducing competitive interactions (MacArthur, 1958).

#### Overyielding

A phenomenon in which greater yield (biomass or productivity) is observed in mixtures than expected statistically based on monocultures of the component species.

**Resistance** Integrated stability, including the resistance to changes in ecosystem properties in response to a stress or disturbance (Helfgott, 2015; Yi & Jackson, 2021).

**Resilience** The ability of ecosystem properties to return to a pre-disturbance condition after a disturbance (Fraccascia et al., 2018).

#### Resource partitioning

Occurs when species use different portions of the available resource pool such that the existing resource pool is more completely used in higher-diversity communities compared with monocultures (Barry et al., 2019).

#### Response and effect traits

Response traits describe a plant’s response to the biotic and abiotic environment, while effect traits describe the effect of a plant on ecosystem functioning (Lavorel & Garnier, 2002; Violle et al., 2007).

**Stability** The capacity of an ecosystem to maintain its structure and function over time, despite disturbances and environmental fluctuations.

**Stress** Adverse abiotic or biotic environmental conditions that exceed an organism’s range of tolerance, potentially harming its growth, survival, or reproduction.

### Measurements

Soils (1 composite of 10 cores/100 m^2^ plot and 20 cores/400 m^2^ plot) from 3 depths (0-15, 15-30, 30-60 cm) were archived at the start of the experiment. All trees are measured annually for tree survival and growth, measured as height, stem basal diameter and diameter at breast height once it is reached. We also collected multispectral imagery and LiDAR point clouds over the experiment. Herbivore and disease monitoring will take place both opportunistically (e.g., as infestations or outbreaks occur) and through periodic (one in ∼five year) stratified random surveys of end-of-season (August/September) generalist herbivory (leaf chewing and skeletonizing intensity), specialist herbivory (gall and leaf miner abundance), and accumulated pathogen load (foliar fungal damage intensity). Following attainment of reproductive maturity for most species (∼10+ years following establishment), we will also conduct regular stratified random surveys of seed output across diversity treatments.

## Results

Mortality rates of trees (Fig. 4) differed among species (DF=11, SS=5.69, F ratio = 39.46, P <0.0001) and with plot size (DF=1, SS=6.16, F ratio=470, P < 0.0001). Across all species, mortality averaged 5% per year in the 100 m^2^ plots and 17% per year in the 400 m^2^ plots (Table 3). Species also differed in their response to plot size as indicated by the significant interaction (DF=11, SS= 2.9, F ratio=20.1, P < 0.0001) when data for both plot sizes were analyzed together (Table S4).

**Figure 4.**
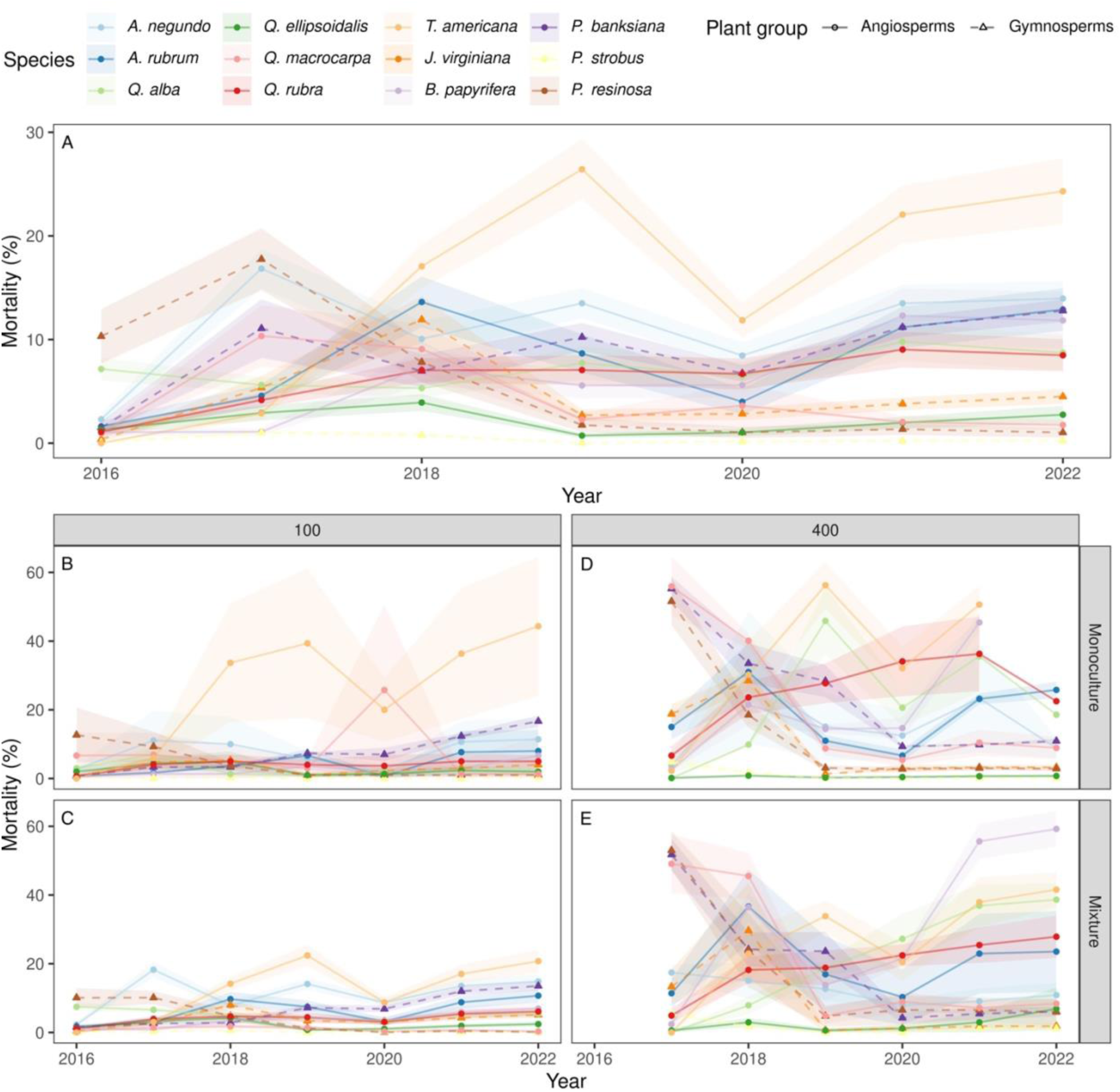
Percent mortality of trees. A) average mortality by species in all plots. Mortality by species in B) 100 m^2^ plots with tree mixtures (any mixture with more than one species), C), 100 m^2^ monocultures, 400 m^2^ mixtures, D) 400 m^2^ monocultures. Mean values per species are shown for each year, with standard error confidence intervals. Mortality percentages were calculated to include any trees newly or previously planted in the plots. The 100 m^2^ plots were planted in 2016, the 400 m^2^ plots were planted in 2017; these were replanted as necessary for three years.

**Table 3.** Average tree mortality rates (%) by species and year across all 100 m^2^ assemblages (top) and 400 m^2^ assemblages (bottom).

|  |  | % mortality |  |  |  |  |  |  |  |  |
| --- | --- | --- | --- | --- | --- | --- | --- | --- | --- | --- |
| 100 m<sup>2</sup> plots |  |  |  |  |  |  |  |  |  |  |
| Species |  | 2016 | 2017 | 2018 | 2019 | 2020 | 2021 | 2022 | Average | SD |
| *Acer negundo* |  | 2.8 | 4.2 | 9.1 | 13.0 | 8.1 | 13.9 | 15.0 | **9.4** | 3.8 |
| *Acer rubrum* |  | 1.7 | 0.5 | 8.3 | 7.6 | 2.4 | 8.5 | 10.0 | **5.6** | 3.5 |
| *Betula papyrifera* |  | 0.8 | 0.3 | 3.0 | 4.3 | 3.1 | 5.5 | 6.8 | **3.4** | 2.1 |
| *Juniperus virginiana* |  | 0.5 | 1.5 | 8.5 | 3.1 | 2.9 | 3.7 | 4.8 | **3.6** | 2.2 |
| *Quercus alba* |  | 6.3 | 2.2 | 4.7 | 4.0 | 3.0 | 4.1 | 6.7 | **4.4** | 1.4 |
| *Quercus ellipsoidalis* |  | 1.5 | 1.3 | 3.6 | 0.8 | 1.0 | 1.6 | 3.9 | **2.0** | 1.2 |
| *Quercus macrocarpa* |  | 2.2 | 2.1 | 2.5 | 1.3 | 0.5 | 0.9 | 6.3 | **2.2** | 1.9 |
| *Quercus rubra* |  | 1.1 | 1.3 | 5.4 | 4.0 | 3.2 | 5.3 | 9.8 | **4.3** | 2.6 |
| *Pinus banksiana* |  | 1.9 | 0.9 | 3.6 | 7.7 | 7.5 | 12.4 | 14.5 | **6.9** | 4.7 |
| *Pinus resinosa* |  | 12.5 | 3.3 | 4.8 | 1.1 | 0.4 | 0.5 | 0.5 | **3.3** | 1.8 |
| *Pinus strobus* |  | 0.2 | 0.0 | 0.9 | 0.0 | 0.2 | 0.2 | 0.2 | **0.2** | 0.3 |
| *Tilia americana* |  | 2.5 | 1.7 | 20.1 | 27.8 | 12.5 | 23.2 | 28.0 | **16.5** | 9.3 |
| **Average** |  | **2.8** | **1.6** | **6.2** | **6.2** | **3.7** | **6.6** | **8.9** | **5.2** |  |
| SD |  | 3.4 | 1.2 | 5.1 | 7.7 | 3.8 | 6.9 | 7.6 | 4.3 |  |
| 400 m<sup>2</sup> plots |  | 2017 | 2018 | 2019 | 2020 | 2021 | 2022 | Average | SD |  |
| *Acer negundo* |  | 7.7 | 27.7 | 14.9 | 12.0 | 21.9 | 9.2 | **15.6** | 7 |  |
| *Acer rubrum* |  | 17.6 | 33.0 | 11.8 | 7.2 | 23.3 | 25.8 | **19.8** | 9 |  |
| *Betula papyrifera* |  | 2.6 | 24.4 | 14.6 | 15.8 | 47.2 | 59.9 | **27.4** | 20 |  |
| *Juniperus virginiana* |  | 19.6 | 29.5 | 1.3 | 2.7 | 3.1 | 3.2 | **9.9** | 11 |  |
| *Quercus alba* |  | 0.1 | 9.6 | 38.1 | 20.7 | 33.7 | 24.9 | **21.2** | 13 |  |
| *Quercus ellipsoidalis* |  | 0.2 | 1.1 | 0.3 | 0.5 | 0.9 | 1.5 | **0.8** | 0 |  |
| *Quercus macrocarpa* |  | 55.1 | 42.8 | 8.8 | 5.9 | 10.0 | 9.2 | **22.0** | 19 |  |
| *Quercus rubra* |  | 5.9 | 22.6 | 22.6 | 29.2 | 31.4 | 24.2 | **22.7** | 8 |  |
| *Pinus banksiana* |  | 55.3 | 33.7 | 27.8 | 8.8 | 9.7 | 10.6 | **24.3** | 17 |  |
| *Pinus resinosa* |  | 51.8 | 19.6 | 3.4 | 3.2 | 3.4 | 3.2 | **14.1** | 18 |  |
| *Pinus strobus* |  | 4.9 | 1.8 | 0.1 | 0.1 | 0.3 | 0.3 | **1.2** | 2 |  |
| *Tilia americana* |  | 1.7 | 29.6 | 40.8 | 23.4 | 41.2 | 42.2 | **29.8** | 14 |  |
| **Average** |  | **18.5** | **23.0** | **15.4** | **10.8** | **18.9** | **17.9** | **17.4** |  |  |
| SD |  | 22.3 | 12.9 | 14.2 | 9.6 | 16.6 | 18.4 | 9.4 |  |  |

Within the 100 m^2^ plots, mortality of trees significantly differed between species (DF=11, SS=4.85, F ratio = 44.45, P <0.0001), and across diversity treatments (DF=1, SS=0.102, F ratio=10.34, P = 0.0013 for PD) with a significant species by diversity interaction (DF=11, SS=0.32, F ratio=2.93, P = 0.0007 for PD) and lower overall mortality in more diverse plots. Species, plot diversity and their interaction significantly predicted mortality, regardless of which diversity metric was used (Table S4). *Tilia americana* and *Acer negundo*, in particular, showed decreasing mortality with increasing diversity of the plots (Fig. 5 A-C). Composition of the plots in conjunction with diversity was also important in predicting mortality. Highest mortality occurred in low diversity angiosperm mixtures (Fig. 5 E-F). Only lower diversity mixtures were composed of angiosperms only. The proportion of angiosperms in a mixture significantly predicted percent tree mortality (DF=1, SS=0.08, F ratio = 9.5, P = 0.0021) when species, phylogenetic diversity (or other metric of diversity) and the interactions with species were included in the model. When phylogenetic diversity (or other metric of diversity) was not included in the model, the proportion of angiosperms was not directly significant in predicting tree mortality but the interaction of species and proportion of angiosperms in the mixture was significant (DF=11, SS=0.295, F ratio=3.1, P=0.0004, Table S4). When only the 400 m^2^ plots were included in the analysis, species significantly differed in mortality (DF=11, SS=3.79, F ratio=39.91, P<0.0001, Fig. 4, Table S4), but no diversity effect was discernable, given these comprise 36 monoculture plots and only five 12-species plots.

**Figure 5.**
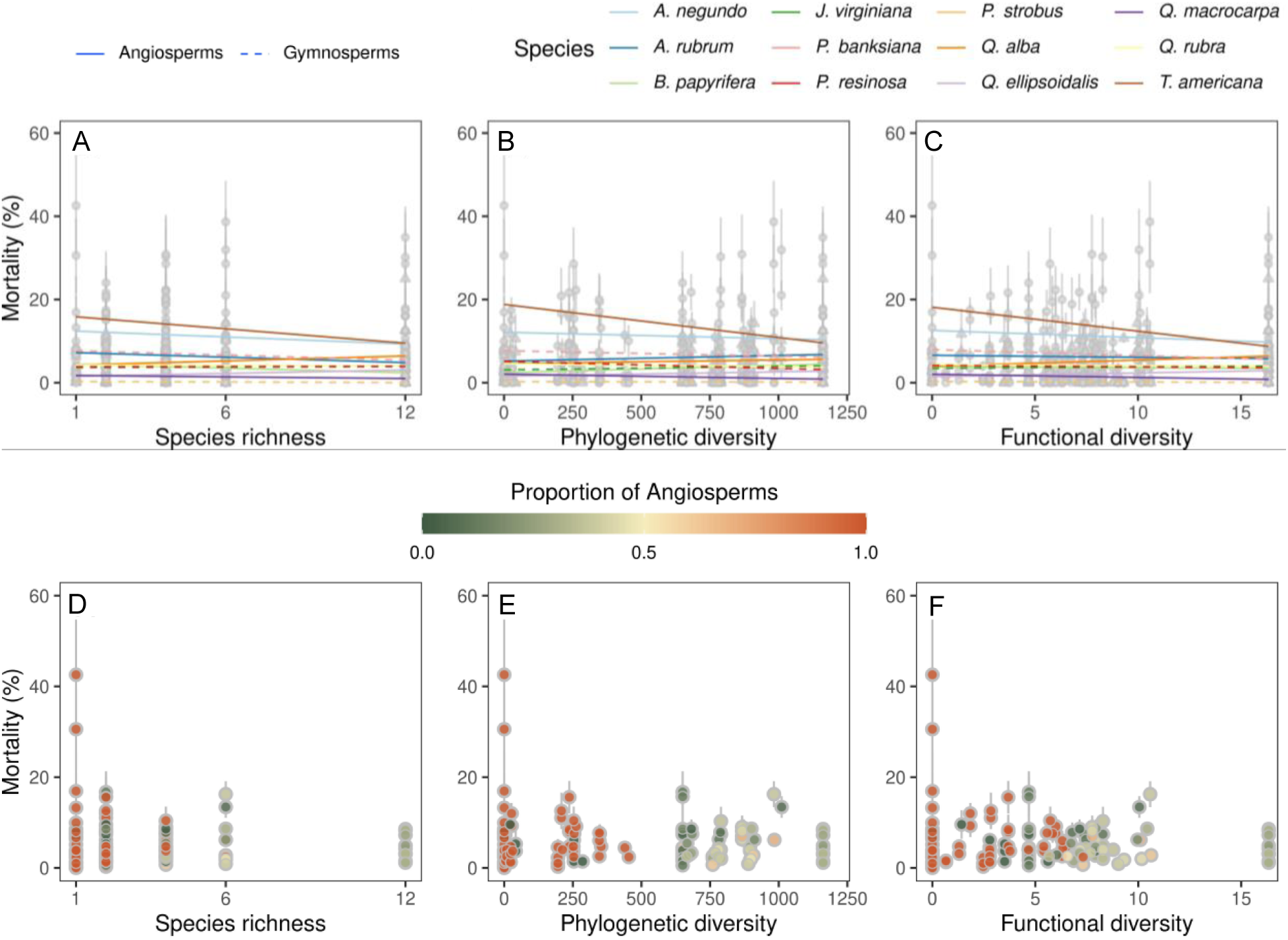
Tree mortality by species or plot composition in relation to plot diversity. In the top three panels, percent mortality per species averaged over time is shown in relation to (A) species richness level, (B) phylogenetic diversity, calculated as the sum of the phylogenetic branch lengths of all species in the assemblage (Faith’s PD) of all species in the assemblage with monocultures shown as zero phylogenetic diversity, and (C) functional diversity calculated as the sum of functional distances for a suite of eight traits, with monocultures shown as zero functional diversity. Linear models are fit to each species, shown as different colored lines. Solid lines are angiosperms and dashed lines are gymnosperms. In the bottom three panels, mean percent mortality by assemblage averaged across time is shown in relation to (D) species richness, (E) phylogenetic diversity and (F) functional diversity. The proportion of angiosperm tree species in each plot is color indicated, with increasing proportion of angiosperms shown as more orange and greater proportion of conifers as more green.

## Discussion

Mortality of trees significantly differed with plot size, species and diversity. Highest mortality occurred in low diversity broadleaf (angiosperm) plots. Despite irrigation, it is likely that multiple years of successive drought caused greater mortality in angiosperm monocultures (Fig. 5), where lack of shading by fast-growing conifer neighbors may have caused excessive evaporative loss. *Tilia americana* had particularly high levels of mortality in both plot sizes, particularly in monocultures and in low diversity angiosperm mixtures, which we hypothesize resulted from its dependence on shade from heterospecific neighbors, given that its growth in FAB1 was enhanced by shading (Kothari et al., 2021). *Acer negundo* showed a similar pattern. These results point to the importance of forest diversity in preventing mortality of vulnerable tree species during early phases of establishment.

The large variation in functional and phylogenetic composition across species richness levels create an important foundation to test critical hypotheses about the consequences of species and lineages and the diversity effects from their interactions for ecosystems and other trophic levels. The lessons that we stand to learn from this major experiment, in concert with similar experiments established globally (Paquette et al. 2018), will provide critical insights that can inform ecosystem management on our rapidly changing planet. Nevertheless, establishing and maintaining a large manipulative tree diversity experiment pose a number of challenges. Designing the experiment faced several issues. The 12 species in the experiment were chosen primarily based on their survival percentage in a preliminary (pre-FAB) planting experiment at Cedar Creek Ecosystem Science Reserve without watering, weeding or fertilizer amendments. With this species pool, it was not possible to create diversity treatments where FV and PV or FD and PD were fully orthogonal. Different metrics and ways of calculating phylogenetic and functional diversity also give different spreads of data. The high mortality during establishment of the experiment required us to replant trees from 2017-2020 each year in the 100 m^2^ plots and from 2018-2022 in the 400 m^2^ plots. In 2022, we had shortfalls in tree availability for replanting the 400 m^2^ plots due to the COVID-19 pandemic. These trees were replaced in spring 2023, the last year of replanting. In 2022, Cedar Creek experienced an extreme summer drought, and some new plantings were especially susceptible to these conditions. Finally, the large size of the experiment makes exhaustive measurements of all individual trees beyond the annual growth survey challenging, and subsampling is critical. We are increasingly relying on remotely sensed measures of trees and plots to complement subsampling on the ground (Fig. 3). The aim of this initial manuscript on FAB2 is to present the experimental design and initial mortality results as well as to clarify the central hypotheses and document the challenges. Future studies in the near- and long-term are planned to test the mechanistic hypotheses presented.

## Acknowledgements

We wish to thank Allison Scott for help with tree mortality data collection and replanting and Christopher Buyarski, Will Dunker, Gloria Perez, Sydney Schiffner, Jason Frey and the Osprey Wilds Environmental Learning Center and the Minnesota Iowa Conservation Corps for assistance sourcing seeds and contributing to in the support of this large experiment. We thank the NSF Cedar Creek Long Term Ecological Research grant DEB: 1831944 for funding establishment of the experiment and the NSF ASCEND BII DBI: 2021898 for funding the remote sensing imagery. Data from FAB are accessible through the DRUM repository: https://conservancy.umn.edu/handle/11299/218359, EcoSIS repository: https://ecosis.org/package/fab-leaf-spectra-across-a-light-gradient-at-cedar-creek-lter, https://ecosis.org/package/leaf-spectra-forest-and-biodiversity-experiment-cedar-creek-lter and Cedar Creek Data Catalog: https://cedarcreek.umn.edu/research/

## Statement of Inclusion

We acknowledge that the FAB2 experiment was established on Dakota and Ojibwe land. Our study brings together contributors from diverse backgrounds representing diversity in gender, age, sexual orientation, race and ethnicity and nationality. All authors were engaged early on with the manuscript preparation to ensure that the diverse sets of perspectives they represent was considered from the onset. The authors all work or have worked at Cedar Creek Ecosystem Science Reserve and are dedicated—through the processes of research, conservation, education and public engagement—to inclusion and bridging the gaps between science, community and government.

## Supplemental Information

**Figure S1.**
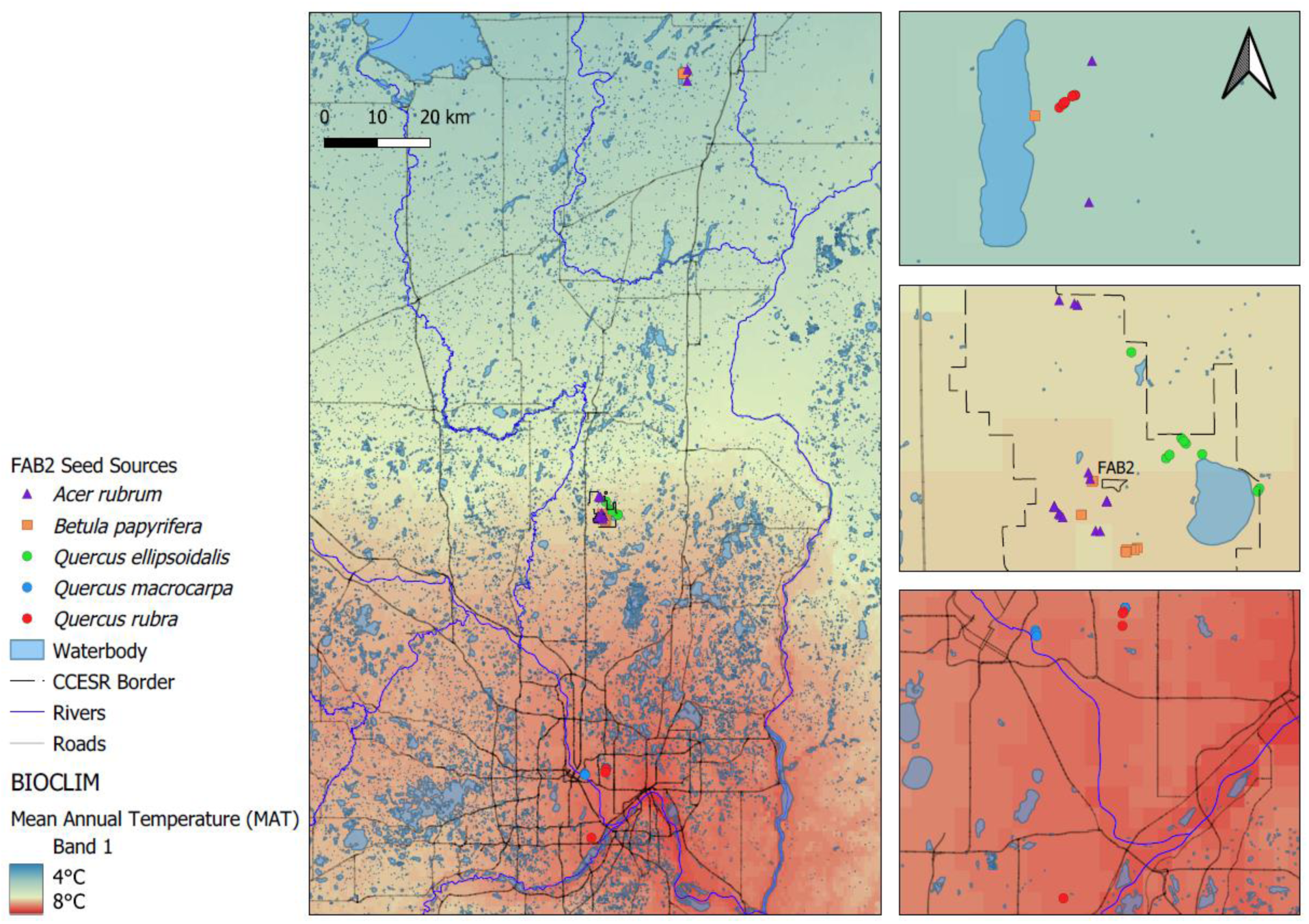
For five species, seeds were collected from known mother trees within 100 km of the Cedar Creek Ecosystem Science Reserve (CCESR), East Bethel, MN, USA. Three primary locations were utilized, representing a 4 °C range in mean annual temperature. Left panel is centered on CCESR (45°25’ N, 93°10’ W). Panels on the right highlight primary collection sites of Sandstone, MN (top), CCESR (middle) and Minneapolis/St Paul, MN (bottom).

**Figure S2.**
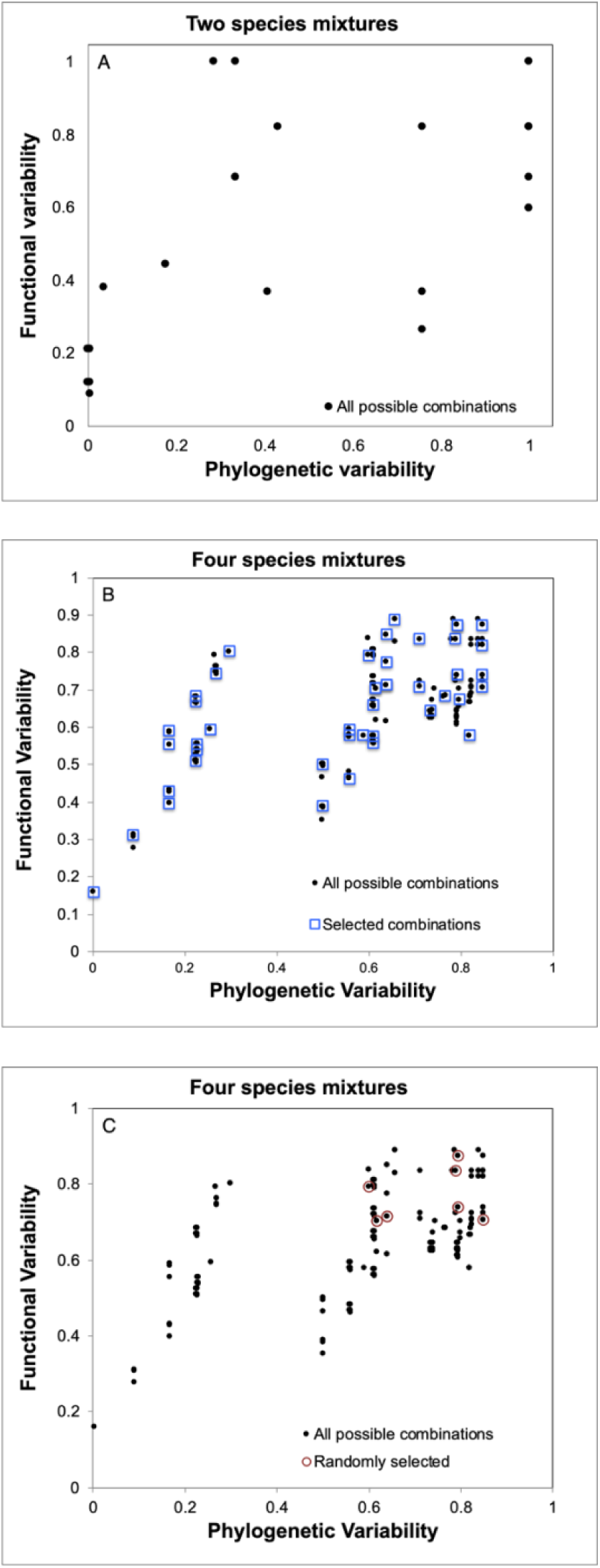
Possible treatment combinations based on functional variability (FV) and phylogenetic variability (PV) of tree species for (A) two species and (B) four species mixtures. For the four species mixtures, the PV-FV selected mixtures are shown in blue squares (B), and the randomly drawn combinations of four species are shown in red circles (C). Random draws tended to have high PV and high FV combinations and to include both angiosperms and gymnosperms. PV and FV vary between 0 and 1 and are independent of species richness (Helmus 2007). See text for details.Figure S4. Original phylogeny from design phase (A) and current phylogeny (B) used for calculating phylogenetic variability and diversity.

**Figure S3.**
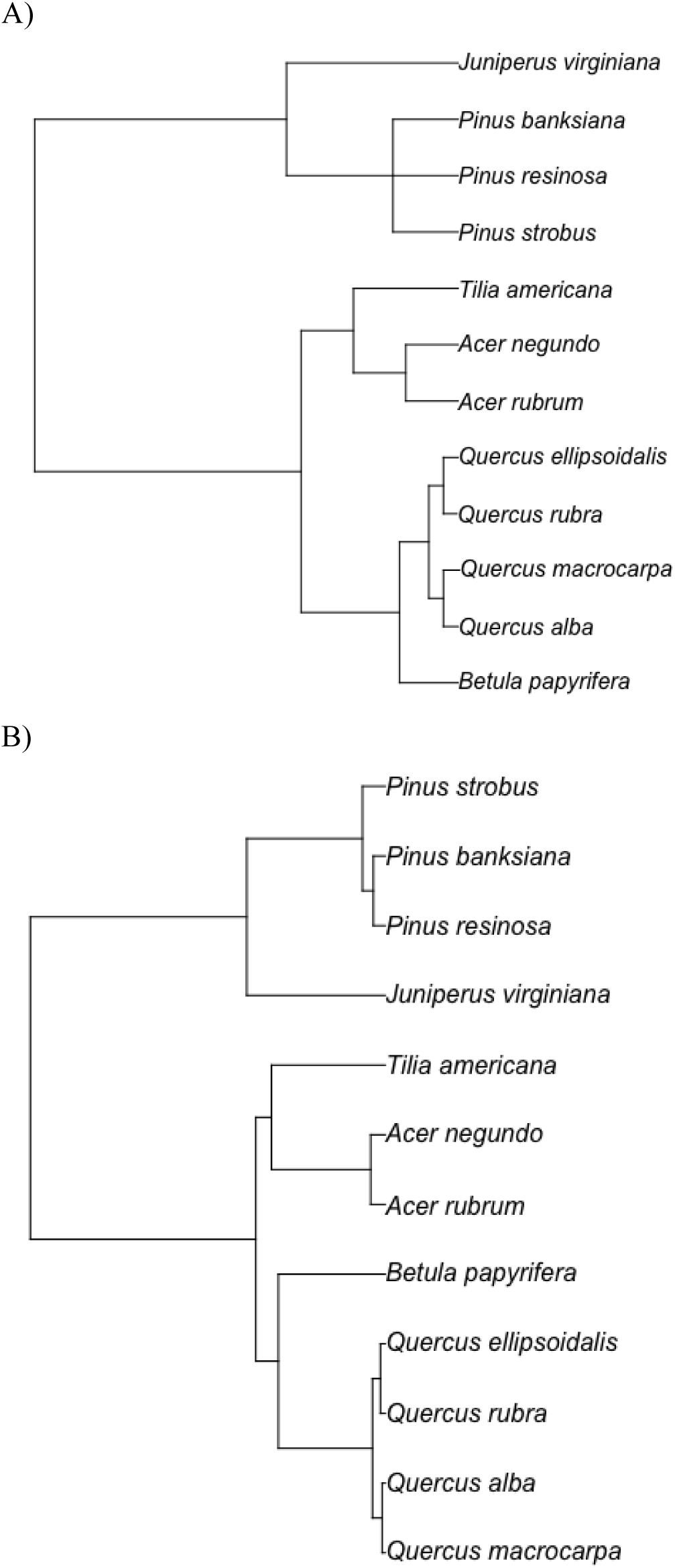
Original phylogeny from design phase (A) and current phylogeny (B) used for calculating phylogenetic variability and diversity.

**Fig. S4.**
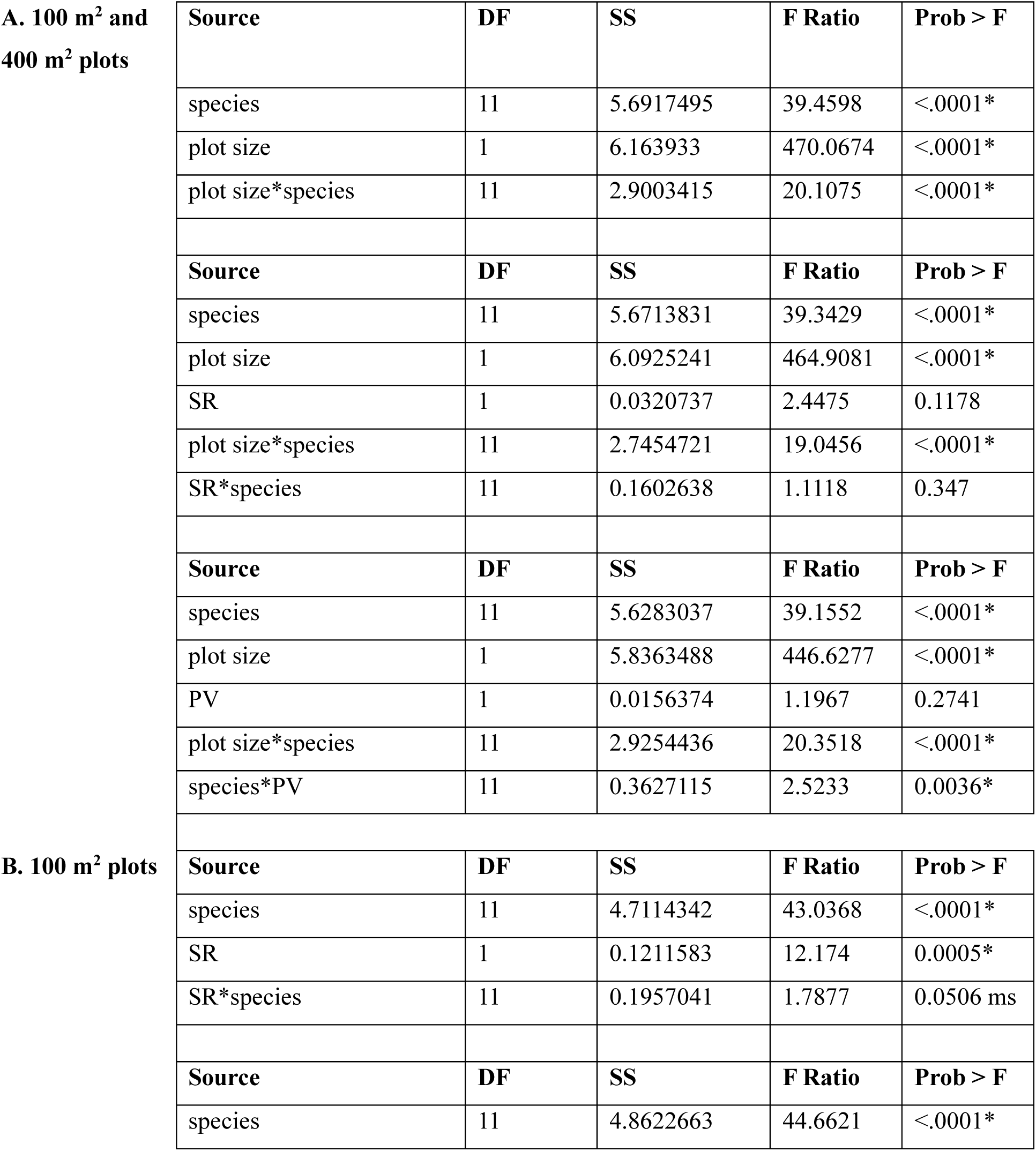
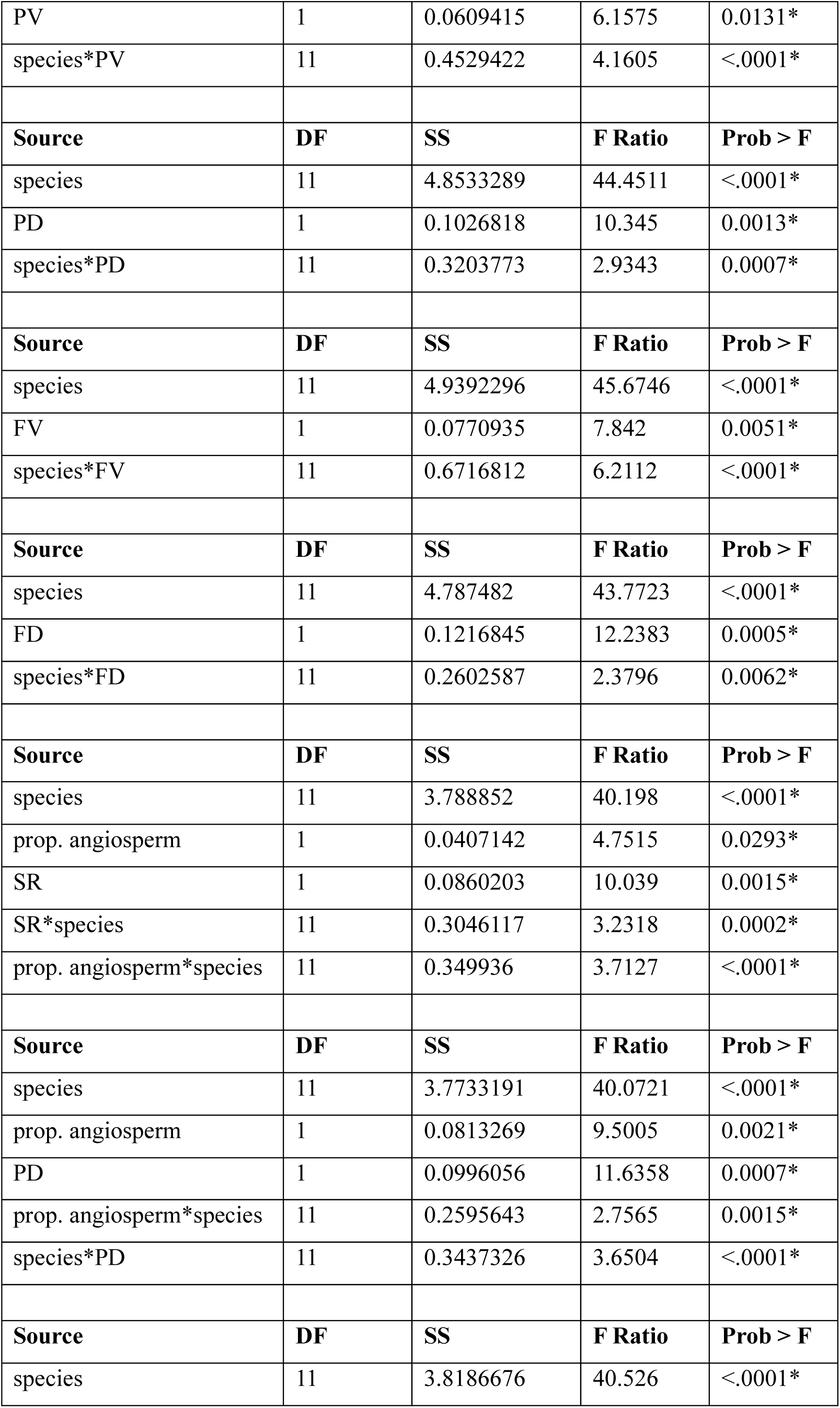
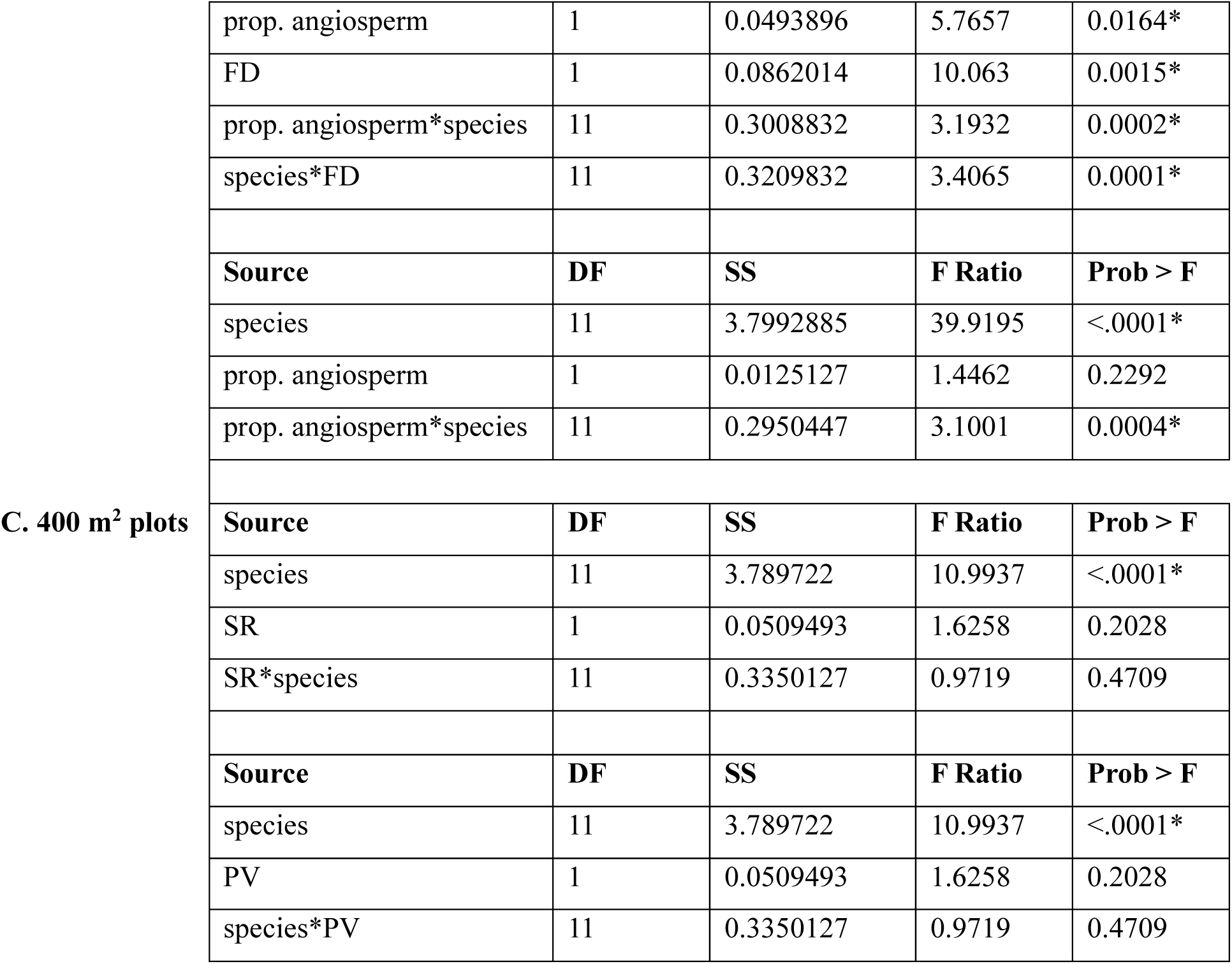
Linear model results. In all analyses, percent mortality (calculated per species in every plot where it was planted) was treated as the independent variable. Predictor variables and their interactions were included as indicated. Diversity metrics were abbreviated as follows: SR = species richness, PV = phylogenetic variability, FV = functional variability, PD = phylogenetic diversity (Faith’s), FD = functional diversity (sum of dendrogram distances). DF = degrees of freedom, SS= sum of squares. See text for details of how the metrics were calculated. In A, data for 100 m^2^ and 400 m^2^ plots were combined to test for a plot size effect. In B, only 100 m^2^ plots were included. In C, only 400 m^2^ plots were included.

## Literature Cited

Averill, C., Bhatnagar, J. M., Dietze, M. C., Pearse, W. D., & Kivlin, S. N. (2019). Global imprint of mycorrhizal fungi on whole-plant nutrient economics. In Proceedings of the National Academy of Sciences (Vol. 116, Issue 46, p. 23163). 10.1073/pnas.1906655116

Barry, K. E., Mommer, L., van Ruijven, J., Wirth, C., Wright, A. J., Bai, Y., Connolly, J., De Deyn, G. B., de Kroon, H., Isbell, F., Milcu, A., Roscher, C., Scherer-Lorenzen, M., Schmid, B., & Weigelt, A. (2019). The future of complementarity: Disentangling causes from consequences. In Trends in Ecology & Evolution (Vol. 34, Issue 2, pp. 167–180). 10.1016/j.tree.2018.10.013

Brooker, R. W., Maestre, F. T., Callaway, R. M., Lortie, C. L., Cavieres, L. A., Kunstler, G., Liancourt, P., Tielbörger, K., Travis, J. M. J., Anthelme, F., Armas, C., Coll, L., Corcket, E., Delzon, S., Forey, E., Kikvidze, Z., Olofsson, J., Pugnaire, F., Quiroz, C. L., … Michalet, R. (2008). Facilitation in plant communities: The past, the present, and the future. In Journal of Ecology (Vol. 96, Issue 1, pp. 18–34). 10.1111/j.1365-2745.2007.01295.x

Bryant, R. L., Kothari, S., Cavender-Bares, J., Curran, S. J., Grossman, J. J., Hobbie, S. E., Montgomery, R., Nash, C., Neumiller, G. C., & See, C. R. (n.d.). Independent effects of tree diversity on above- and belowground carbon pools.

Cadotte, M., Cavender-Bares, J., Oakley, T., & Tilman, D. (2009). Using phylogenetic, functional and trait diversity to understand patterns of plant community productivity. In PLoS ONE (Vol. 4, Issue 5, p. e5695). doi:10.1371/journal.pone.0005695

Cadotte, M. W., Cardenale, B. J., & Oakley, T. H. (2008). Evolutionary history and the effect of biodiversity on plant productivity. In Proceedings of the National Academy of Sciences of the United States of America (Vol. 105, pp. 17012–17017).

Cadotte, M. W., & Dinnage, R. (2012). Phylogenetic diversity promotes ecosystem stability. In Ecology (Vol. 93, pp. S223–S233).

Castagneyrol, B., B. Giffard, C. Péré, and H. Jactel.(2013). Plant Apparency, an Overlooked Driver of Associational Resistance to Insect Herbivory. Journal of Ecology 101(2) 418–29.

Cavender-Bares, J., Kozak, K. H., Fine, P. V. A., & Kembel, S. W. (2009). The merging of community ecology and phylogenetic biology. Ecology Letters, 12(7), 693–715. 10.1111/j.1461-0248.2009.01314.x

Cavender-Bares, J., and P. B. Reich. 2012. Shocks to the system: Community assembly of the oak savanna in a 40-year fire frequency experiment. Ecology 93:S52–S69.

Cavender-Bares, J. M., Schweiger, A. K., Gamon, J. A., Gholizadeh, H., Helzer, K., Lapadat, C., Madritch, M. D., Townsend, P. A., Wang, Z., & Hobbie, S. E. (2021). Remotely detected aboveground plant function predicts belowground processes in two prairie diversity experiments. In Ecological Monographs: Vol. DOI:10.1002/ecm.1488. DOI:10.1002/ecm.1488

Cavender-Bares, J., Schneider, F. D., Santos, M. J., Armstrong, A., Carnaval, A., Dahlin, K. M., Fatoyinbo, L., Hurtt, G. C., Schimel, D., & Townsend, P. A. (2022). Integrating remote sensing with ecology and evolution to advance biodiversity conservation. In Nature Ecology & Evolution (Vol. 6, Issue 5, pp. 506–519).

Cavender-Bares, J., Schweiger, A. K., Pinto-Ledezma, J. N., & Meireles, J. E. (2020). Applying Remote Sensing to Biodiversity Science. In J. Cavender-Bares, J. A. Gamon, & P. A. Townsend (Eds.), Remote Sensing of Plant Biodiversity (pp. 13–42). Springer International Publishing. 10.1007/978-3-030-33157-3_2

Cavender-Bares, J., Townsend, P. A., & Gamon, J. A. (2020). Remote Sensing of Plant Biodiversity. Springer Nature. 10.1007/978-3-030-33157-3

Chapin, F. S., Matson, P. A., & Vitousek, P. M. (2011). Species Effects on Ecosystem Processes. In F. S. Chapin, P. A. Matson, & P. M. Vitousek (Eds.), Principles of Terrestrial Ecosystem Ecology (pp. 321–336). Springer New York. 10.1007/978-1-4419-9504-9_11

Cline, L. C., Hobbie, S. E., Madritch, M. D., Buyarski, C. R., Tilman, D., & Cavender-Bares, J. M. (2018). Resource availability underlies the plant-fungal diversity relationship in a grassland ecosystem. In Ecology (Vol. 99, Issue 1, pp. 204–216). 10.1002/ecy.2075

Crawford, M. S., Barry, K. E., Clark, A. T., Farrior, C. E., Hines, J., Ladouceur, E., Lichstein, J. W., Maréchaux, I., May, F., Mori, A. S., Reineking, B., Turnbull, L. A., Wirth, C., & Rüger, N. (2021). The function-dominance correlation drives the direction and strength of biodiversity– ecosystem functioning relationships. Ecology Letters, 24(9), 1762–1775. 10.1111/ele.13776

Dahlin, K. M. (2016). Spectral diversity area relationships for assessing biodiversity in a wildland– agriculture matrix. In Ecological Applications (Vol. 26, Issue 8, pp. 2758–2768). 10.1002/eap.1390

Damien, M., Jactel, H., Meredieu, C., Régolini, M., van Halder, I., & Castagneyrol, B. (2016). Pest damage in mixed forests: Disentangling the effects of neighbor identity, host density and host apparency at different spatial scales. In Forest Ecology and Management (Vol. 378, pp. 103–110). 10.1016/j.foreco.2016.07.025

Eisenhauer, N., Bonkowski, M., Brose, U., Buscot, F., Durka, W., Ebeling, A., Fischer, M., Gleixner, G., Heintz-Buschart, A., Hines, J., Jesch, A., Lange, M., Meyer, S., Roscher, C., Scheu, S., Schielzeth, H., Schloter, M., Schulz, S., Unsicker, S., … Schmid, B. (2019). Biotic interactions, community assembly, and eco-evolutionary dynamics as drivers of long-term biodiversity– ecosystem functioning relationships. In Research Ideas and Outcomes (Vol. 5, p. e47042).

Endara, M.-J., & Coley, P. D. (2011). The resource availability hypothesis revisited: A meta-analysis. In Functional Ecology (Vol. 25, Issue 2, pp. 389–398). 10.1111/j.1365-2435.2010.01803.x

Ewel, J. J., Celis, G., & Schreeg., L. (2015). Steeply increasing growth differential between mixture and monocultures of tropical trees. In Biotropica (Vol. 47, pp. 162–171).

Faith, D. P. (1992). Conservation evaluation and phylogenetic diversity. In Biol. Conserv. (Vol. 61, pp. 1–10).

Fernández-Tschieder, E., & Binkley, D. (2018). Linking competition with Growth Dominance and production ecology. Forest Ecology and Management, 414, 99–107. 10.1016/j.foreco.2018.01.052

Flynn, D. F. B., Mirotchnick, N., Jain, M., Palmer, M. I., & Naeem, S. (2011). Functional and phylogenetic diversity as predictors of biodiversity–ecosystem-function relationships. In Ecology (Vol. 92, Issue 8, pp. 1573–1581). 10.1890/10-1245.1

Fraccascia, L., Giannoccaro, I., & Albino, V. (2018). Resilience of complex systems: State of the art and directions for future research. In Complexity (Vol. 2018, p. 3421529). 10.1155/2018/3421529

Gonzalez, A., Vihervaara, P., Balvanera, P., Bates, A. E., Bayraktarov, E., Bellingham, P. J., Bruder, A., Campbell, J., Catchen, M. D., Cavender-Bares, J., Chase, J., Coops, N., Costello, M. J., Dornelas, M., Dubois, G., Duffy, E. J., Eggermont, H., Fernandez, N., Ferrier, S., … Wright, E. (2023). A global biodiversity observing system to unite monitoring and guide action. Nature Ecology & Evolution. 10.1038/s41559-023-02171-0

Grossman, J. J., Butterfield, A. J., Cavender-Bares, J., Hobbie, S. E., Reich, P. B., Gutknecht, J., & Kennedy, P. G. (2019). Non-symbiotic soil microbes are more strongly influenced by altered tree biodiversity than arbuscular mycorrhizal fungi during initial forest establishment. In FEMS Microbiology Ecology (Vol. 95, Issue 10). 10.1093/femsec/fiz134

Grossman, J. J., Cavender-Bares, J., & Hobbie, S. E. (2020). Functional diversity of leaf litter mixtures slows decomposition of labile but not recalcitrant carbon over two years. In Ecological Monographs (Vol. 90, Issue 3, p. e01407). 10.1002/ecm.1407

Grossman, J. J., Cavender-Bares, J., Hobbie, S. E., Reich, P. B., & Montgomery, R. A. (2017). Species richness and traits predict overyielding in stem growth in an early-successional tree diversity experiment. In Ecology (Vol. 98, Issue 10, pp. 2601–2614). doi:10.1002/ecy.1958

Grossman, J. J., Vanhellemont, M., Barsoum, N., Bauhus, J., Bruelheide, H., Castagneyrol, B., Cavender-Bares, J., Eisenhauer, N., Ferlian, O., Gravel, D., Hector, A., Jactel, H., Kreft, H., Mereu, S., Messier, C., Muys, B., Nock, C., Paquette, A., Parker, J., … Verheyen, K. (2018). Synthesis and future research directions linking tree diversity to growth, survival, and damage in a global network of tree diversity experiments. In Environmental and Experimental Botany (Vol. 152, pp. 68–89). 10.1016/j.envexpbot.2017.12.015

Haggar, J. P., & Ewel, J. J. (1997). Primary productivity and resource partitioning in model tropical ecosystems. In Ecology (Vol. 78, pp. 1211–1221).

Hector, A., B. Schmid, C. Beierkuhnlein, M. C. Caldeira, M. Diemer, P. G. Dimitrakopoulos, J. A. Finn, H. Freitas, P. S. Giller, J. Good, R. Harris, P. Hogberg, K. Huss-Danell, J. Joshi, A. Jumpponen, C. Korner, P. W. Leadley, M. Loreau, A. Minns, C. P. H. Mulder, G. O’Donovan, S. J. Otway, J. S. Pereira, A. Prinz, D. J. Read, M. Scherer-Lorenzen, E. D. Schulze, A. S. D. Siamantziouras, E. M. Spehn, A. C. Terry, A. Y. Troumbis, F. I. Woodward, S. Yachi, and J. H. Lawton. 1999. Plant diversity and productivity experiments in European grasslands. Science 286:1123–1127.

Helfgott, A. E. R. (2015). Operationalizing resilience: Conceptual, mathematical and participatory frameworks for understanding, measuring and managing resilience [Thesis].

Helmus, M. R. (2007). Phylogenetic measures of biodiversity. In American Naturalist (Vol. 169, Issue 3, pp. E68–E83).

Hobbie, S. E. (1994). Plant species’ effects are larger than those of increased temperature on microbial activity in tussock tundra litter-soil microcosms. In Bulletin of the Ecological Society of America (Vol. 75, Issue 2, p. 95).

Hobbie, S. E. (1995). Direct and indirect effects of plant species on biogeochemical processes in arctic ecosystems. In I. Chapin F. S. & Ch. Körner (Eds.), Arctic and alpine biodiversity: Patterns, causes and ecosystem consequences (pp. 213–224). Springer-Verlag.

Hobbie, S. E. (2015). Plant species effects on nutrient cycling: Revisiting litter feedbacks. In Trends in ecology & evolution (Vol. 30, Issue 6, pp. 357–363).

Hobbie, S., Reich, P., Oleksyn, J., Ogdahl, M., Zytkowiak, R., Hale, C., & Karolewski, P. (2006). Tree species effects on decomposition and forest floor dynamics in a common garden. In Ecology (Vol. 87, Issue 9, pp. 2288–2297).

Hooper, D. U., Chapin, F. S., Ewel, J. J., Hector, A., Inchausti, P., Lavorel, S., Lawton, J. H., Lodge, D. M., Loreau, M., Naeem, S., Schmid, B., Setälä, H., Symstad, A. J., Vandermeer, J., & Wardle, D. A. (2005). Effects of Biodiversity on Ecosystem Functioning: A Consensus of Current Knowledge. In Ecological Monographs (Vol. 75, Issue 1, pp. 3–35). 10.1890/04-0922

Hooper, D. U., & Dukes, J. S. (2004). Overyielding among plant functional groups in a long-term experiment. In Ecology Letters (Vol. 7, Issue 2, pp. 95–105).

Hutchison, C., Gravel, D., Guichard, F., & Potvin, C. (2018). Effect of diversity on growth, mortality, and loss of resilience to extreme climate events in a tropical planted forest experiment. In Scientific Reports (Vol. 8, Issue 1, p. 15443). 10.1038/s41598-018-33670-x

Jenkins, J. C., Chojnacky, D. C., Heath, L. S., & Birdsey, R. A. (2003). National scale biomass estimators for United States tree species. In Forest Science (Vol. 49, pp. 12–35).

Keesing, F., & Ostfeld, R. S. (2021). Dilution effects in disease ecology. Ecology Letters, 24(11), 2490– 2505. 10.1111/ele.13875

Kembel, S. W., Ackerly, D. D., Blomberg, S. P., Cornwell, W. K., Cowan, P. D., Helmus, M. R., Morlon, H., & Webb, C. O. (2010). Picante: R tools for integrating phylogenies and ecology. In Bioinformatics (Issue 26, pp. 1463–1464).

Kothari, S., Montgomery, R. A., & Cavender-Bares, J. (2021). Physiological responses to light explain competition and facilitation in a tree diversity experiment. In Journal of Ecology (Vol. 109, Issue 5, pp. 2000–2018). 10.1111/1365-2745.13637

Ladegaard-Pedersen, P., B. Elberling, and L. Vesterdal. 2005. Soil carbon stocks, mineralization rates, and CO2 effluxes under 10 tree species on contrasting soil types. Canadian Journal of Forest Research 35:1277–1284.

Larkin, Daniel J., Glasenhardt, Mary-Claire, Williams, Evelyn W., Karimi, Nisa, Barak, Rebecca S., Leavens, Emma, and Hipp, Andrew L.. 2023. “ Evolutionary History Shapes Grassland Productivity through Opposing Effects on Complementarity and Selection.” Ecology 104(8): e4129. 10.1002/ecy.4129

Lavorel, S., & Garnier, E. (2002). Predicting changes in community composition and ecosystem functioning from plant traits: Revisiting the Holy Grail. In Functional Ecology (Vol. 16, Issue 16, pp. 545–556).

Lavorel, S., Grigulis, K., McIntyre, S., Williams, N. S. G., Garden, D., Dorrough, J., Berman, S., Quétier, F., Thébault, A., & Bonis, A. (2008). Assessing functional diversity in the field – methodology matters! In Functional Ecology (Vol. 22, Issue 1, pp. 134–147). 10.1111/j.1365-2435.2007.01339.x

Leadley, P., Archer, E., Bendandi, B., Cavender-Bares, J., Davalos, L., DeClerck, F., Gann, G. D., Gonzales, E. K., Krug, C. B., Metzger, J. P., Nicholson, E., Niinemets, Ü., Obura, D., Strassburg, B., Tansey, B., Verburg, P. H., Vidal, A., Watson, J. E. M., Woodley, S., & Yasuhara, M. (2022). Setting ambitious international restoration objectives for terrestrial ecosystems for 2030 and beyond. In PLOS Sustainability and Transformation (Vol. 1, Issue 12, p. e0000039). 10.1371/journal.pstr.0000039

Loreau, M., & Hector, A. (2001). Partitioning selection and complementarity in biodiversity experiments. In Nature (Vol. 412, p. 72). 10.1038/35083573

MacArthur, R. H. (1958). Population ecology of some warblers of northeastern coniferous forests. In Ecology (Vol. 39, Issue 4, pp. 599–619).

Maillard, F., Beatty, B., Park, M., Adamczyk, S., Adamczyk, B., Cavender-Bares, J., Hobbie, S. E., & Kennedy, P. G. (2023). Soil microbial community structure predicts fungal necromass decomposition rates across diverse temperate forest types. In Soil Biology and Biochemistry (Vol. SBB20767, in review).

Messier, C., Bauhus, J., Sousa-Silva, R., Auge, H., Baeten, L., Barsoum, N., Bruelheide, H., Caldwell, B., Cavender-Bares, J., & Dhiedt, E. (2021). For the sake of resilience and multifunctionality, let’s diversify planted forests! In Conservation Letters (p. e12829).

Mouchet, M. A., Villéger, S., Mason, N. W., & Mouillot, D. (2010). Functional diversity measures: An overview of their redundancy and their ability to discriminate community assembly rules. In Functional Ecology (Vol. 24, Issue 4, pp. 867–876).

Muiruri, E. W., Barantal, S., Iason, G. R., Salminen, J.-P., Perez-Fernandez, E., & Koricheva, J. (2019). Forest diversity effects on insect herbivores: Do leaf traits matter? In New Phytologist (Vol. 221, Issue 4, pp. 2250–2260). 10.1111/nph.15558

Niinemets, Ü., & Valladares, F. (2006). Tolerance to shade, drought, and waterlogging of temperate Northern Hemisphere trees and shrubs. In Ecological monographs (Vol. 76, Issue 4, pp. 521– 547).

Paquette, A., Hector, A., Castagneyrol, B., Vanhellemont, M., Koricheva, J., Scherer-Lorenzen, M., Verheyen, K., Abdala-Roberts, L., Auge, H., Barsoum, N., Bauhus, J., Baum, C., Bruelheide, H., Castagneyrol, B., Cavender-Bares, J., Eisenhauer, N., Ferlian, O., Ganade, G., Godbold, D., … TreeDivNet. (2018). A million and more trees for science. In Nature Ecology & Evolution (Vol. 2, Issue 5, pp. 763–766). 10.1038/s41559-018-0544-0

Phillips, R. P., Brzostek, E., & Midgley, M. G. (2013). The mycorrhizal-associated nutrient economy: A new framework for predicting carbon–nutrient couplings in temperate forests. In New Phytologist (Vol. 199, Issue 1, pp. 41–51). 10.1111/nph.12221

Reich, P. B., Oleksyn, J., Modrzynski, J., Mrozinski, P., Hobbie, S. E., Eissenstat, D. M., Chorover, J., Chadwick, O. A., Hale, C. M., & Tjoelker, M. G. (2005). Linking litter calcium, earthworms and soil properties: A common garden test with 14 tree species. In Ecology letters (Vol. 8, Issue 8, pp. 811–818).

Revell, L. J. (2012). phytools: An R package for phylogenetic comparative biology (and other things). In Methods in Ecology and Evolution (Vol. 3, Issue 2, pp. 217–223). 10.1111/j.2041-210X.2011.00169.x

Root, R. B. (1973). Organization of a Plant-Arthropod Association in Simple and Diverse Habitats: The Fauna of Collards (Brassica Oleracea). Ecological Monographs, 43(1), 95–124. 10.2307/1942161

Russell, A. E., Cambardella, C. A., Ewel, J. J., & Parkin, T. B. (2004). Species, rotation-frequency, and life-form diversity effects on soil carbon in experimental tropical ecosystems. In Ecological Applications (Vol. 14, pp. 47–60).

Schweiger, A. K., Cavender-Bares, J., Kothari, S., Townsend, P. A., Madritch, M. D., Grossman, J. J., Gholizadeh, H., Wang, & R. Gamon, J. A. (2021). Schweiger, A. K., J. Cavender-Bares, S. Kothari, P. A. Townsend, M. D. Madritch, J. J. Grossman, H. Gholizadeh, R. Wang, and J. A. Gamon. 2021. Coupling spectral and resource-use complementarity in experimental grassland and forest communities. Proce. In Proceedings of the Royal Society B: Biological Sciences (Vol. 288, p. 20211290).

Schwinning, S., & Weiner, J. (1998). Mechanisms determining the degree of size asymmetry in competition among plants. Oecologia, 113(4), 447–455. 10.1007/s004420050397

Sendall, K. M., & Reich, P. B. (2013). Variation in leaf and twig CO2 flux as a function of plant size: A comparison of seedlings, saplings and trees. In Tree Physiology (Vol. 33, Issue 7, pp. 713–729). 10.1093/treephys/tpt048

Simberloff, D. (1970). Taxonomic diversity of island biotas. In Evolution (Vol. 24, pp. 23–47).

Smith, S. A., & Brown, J. W. (2018). Constructing a broadly inclusive seed plant phylogeny. In American Journal of Botany. 10.1002/ajb2.1019

Tahvanainen, J. O., & Root, R. B. (1972). The influence of vegetational diversity on the population ecology of a specialized herbivore, Phyllotreta cruciferae (Coleoptera: Chrysomelidae). Oecologia, 10(4), 321–346. 10.1007/BF00345736

Tilman, D. (1993). Species richness of experimental productivity gradients—How important is colonization limitation? In Ecology (Vol. 74, Issue 8, pp. 2179–2191). 10.2307/1939572

Tilman, D. (1994). Competition and biodiversity in spatially structured habitats. In Ecology (Vol. 75, Issue 1, pp. 2–16).

Tobner, C. M., Paquette, A., Gravel, D., Reich, P. B., Williams, L. J., & Messier, C. (2016). Functional identity is the main driver of diversity effects in young tree communities. In Ecology Letters (Vol. 19, Issue 6, pp. 638–647). 10.1111/ele.12600

Vesterdal, L., I. K. Schmidt, I. Callesen, L. O. Nilsson, and P. Gundersen. 2008. Carbon and nitrogen in forest floor and mineral soil under six common European tree species. Forest Ecology and Management 255:35–48.

Vesterdal, L., N. Clarke, B. D. Sigurdsson, and P. Gundersen. 2013. Do tree species influence soil carbon stocks in temperate and boreal forests? Forest Ecology and Management 309:4–18.

Verheyen, K., Vanhellemont, M., Auge, H., Baeten, L., Baraloto, C., Barsoum, N., Bilodeau-Gauthier, S., Bruelheide, H., Castagneyrol, B., Godbold, D., Haase, J., Hector, A., Jactel, H., Koricheva, J., Loreau, M., Mereu, S., Messier, C., Muys, B., Nolet, P., … Scherer-Lorenzen, M. (2016). Contributions of a global network of tree diversity experiments to sustainable forest plantations. In Ambio (Vol. 45, Issue 1, pp. 29–41). 10.1007/s13280-015-0685-1

Villeger, S., Mason, N. W. H., & Mouillot, D. (2008). New multidimensional functional diversity indices for a multifaceted framework in functional ecology. In Ecology (Vol. 89, Issue 8, pp. 2290–2301). 10.1890/07-1206.1

Violle, C., Navas, M.-L., Vile, D., Kazakou, E., Fortunel, C., Hummel, I., & Garnier, E. (2007). Let the concept of trait be functional! In Oikos (Vol. 116, Issue 5, pp. 882–892). 10.1111/j.2007.0030-1299.15559.x

Webb, C. O., Losos, J. B., & Agrawal, A. A. (2006). Integrating phylogenies into community ecology. In Ecology (Vol. 87, Issue 7, pp. S1–S2).

Wedin, D. A., and D. Tilman. 1990. Species effects on nitrogen cycling: a test with perennial grasses. Oecologia 84:433–441.

White, J. C., Gómez, C., Wulder, M. A., & Coops, N. C. (2010). Characterizing temperate forest structural and spectral diversity with Hyperion EO-1 data. In Remote Sensing of Environment (Vol. 114, Issue 7, pp. 1576–1589). 10.1016/j.rse.2010.02.012

Williams, L. J., Cavender-Bares, J., Townsend, P. A., Couture, J. J., Wang, Z., Stefanski, A., Messier, C., & Reich, P. B. (2021). Remote spectral detection of biodiversity effects on forest biomass. In Nature Ecology & Evolution (Vol. 5, pp. 46–54). 10.1038/s41559-020-01329-4

Williams, L. J., Paquette, A., Cavender-Bares, J., Messier, C., & Reich, P. B. (2017). Spatial complementarity in tree crowns explains overyielding in species mixtures. In Nature Ecology & Evolution (Vol. 1, Issue 4). 10.1038/s41559-016-0063

Wright, A. J., Wardle, D. A., Callaway, R., & Gaxiola, A. (2017). The Overlooked Role of Facilitation in Biodiversity Experiments. In Trends in Ecology & Evolution (Vol. 32, Issue 5, pp. 383–390). 10.1016/j.tree.2017.02.011

Wright, I. J., Reich, P. B., Westoby, M., Ackerly, D. D., Baruch, Z., Bongers, F., Cavender-Bares, J., Chapin, T., Cornelissen, J. H., & Diemer, M. (2004). The worldwide leaf economics spectrum. In Nature (Vol. 428, Issue 6985, pp. 821–827). 10.1038/nature02403

Yi, C., & Jackson, N. (2021). A review of measuring ecosystem resilience to disturbance. In Environmental Research Letters (Vol. 16, Issue 5, p. 053008). 10.1088/1748-9326/abdf09

Zanne, A. E., Tank, D. C., Cornwell, W. K., Eastman, J. M., Smith, S. A., FitzJohn, R. G., McGlinn, D. J., O’Meara, B. C., Moles, A. T., Reich, P. B., Royer, D. L., Soltis, D. E., Stevens, P. F., Westoby, M., Wright, I. J., Aarssen, L., Bertin, R. I., Calaminus, A., Govaerts, R., … Beaulieu, J. M. (2014). Three keys to the radiation of angiosperms into freezing environments. In Nature (Vol. 506, Issue 7486, pp. 89–92). 10.1038/nature12872

